# VesicleVoyager: In vivo selection of surface displayed proteins that direct extracellular vesicles to tissue-specific targets

**DOI:** 10.1101/2025.06.04.657858

**Authors:** Yuki Kawai-Harada, Mehrsa Mardikoraem, Ashley V. Makela, Katherine Lauro, Jeannie Lam, Christopher H. Contag, Daniel Woldring, Masako Harada

## Abstract

The development of technologies for screening proteins that bind to specific tissues in vivo and facilitate delivery of large cargos remains challenging, with most approaches limited to cell culture systems that often yield clinically irrelevant hits. To overcome this limitation, we developed a novel molecular screening platform using an extracellular vesicle (EV) display library. EVs are natural molecular carriers capable of delivering diverse cargos, which can be engineered to enhance specificity and targeting through surface modifications. We constructed an EV-display library presenting monobody repertoires on EV surfaces, with genetic cargo inside the EVs corresponding to the displayed proteins. These libraries were screened for tissue specific delivery through serial passage in mice via sequential intravenous administration in and recovery of tissue-selected EVs and amplification of their encapsulated monobody genes at each passage. Our results demonstrated successful selection of tissue-specific targeting proteins, as revealed by fluorescence and bioluminescence imaging followed by DNA sequencing. To understand the stochastic relationship between displayed proteins and packaged genes, we developed a Markov chain model that quantified selection dynamics and predicted enrichment patterns despite the imperfect correlation between phenotype and genotype. This EV-based monobody screening approach, combined with mathematical modeling, is a significant advancement in targeted drug delivery by leveraging the natural capabilities of EVs with the selection of targeting proteins in a physiologically relevant environment.

## Introduction

The ability to precisely target therapeutic cargo to specific tissues and cells remains one of the greatest challenges in drug delivery. Despite decades of research, less than 2% of systemically administered drugs are estimated to reach their intended targets, highlighting the critical need for innovative targeting solutions.^1^ While numerous approaches exist, we present a platform that leverages the intrinsic properties of extracellular vesicles (EVs) to create an alternative to traditional display technologies that is performed in vivo. We sought to develop our VesicleVoyager system as an in vivo selection strategy to identify targeting molecules in the context of living organ systems with an intact circulatory system providing the selection pressure. As such, targeting molecules are screened in the native context in which they will be used for drug delivery. This is a fundamental shift from conventional screening methods and addresses the critical limitations of other approaches through the strategic integration of extracellular vesicle biology and directed evolution.

Small membrane-bound EVs represent nature’s own delivery system that is capable of transporting bioactive molecules between cells in vivo and delivering functional cargo.^2^ These natural nanoparticles shuttle proteins, nucleic acids, and lipids between cells, serving as critical mediators of intercellular communication.^3^ The molecular signature of EVs have been used as a diagnostic indicators, and EVs have been manipulated for the purpose of developing therapeutic delivery tools.^4,5^ Beyond their diagnostic potential as molecular fingerprints of health and disease, EVs offer the potential of transformative therapeutic delivery capabilities that synthetic alternatives cannot, at this time, match.^4,5^

EV engineering has largely involved the addition of single, or small numbers of, surface modifications and the addition of therapeutic cargo.^6^ Importantly, we have demonstrated that DNA transfection of cells produces EVs that display plasmid-encoded surface proteins and contain the exogenously introduced plasmid DNA (pDNA) within the vesicle. ^7–9^ Much like EV encapsulated RNA, the encapsulated DNA is protected from degradation, which is a critical feature for effective delivery vehicles.^7–9^ This was demonstrated with size exclusion chromatography in which the plasmid DNA and EVs were in the same fraction, and that the DNA in this fraction was protected from DNases.^9^ The size of supercoiled plasmid DNA has been proposed to be smaller than the size of EVs;^10–13^ this and the DNase protection data indicates encapsulation of nucleic acids. These findings lead to the possibility that EV-display libraries could be created, similar to phage display libraries, with the distinct advantages of being compatible with mammalian systems and being screened in vivo with functional readouts.

Phage display technology, pioneered by George P. Smith over three decades ago, remains a cornerstone in discovering biological knowledge and developing pharmaceutical interventions.^14–16^ While alternative screening platforms (such as yeast display, mammalian cell display, bacterial display, and ribosome display) have emerged, phage display remains the most widely used method for screening antibodies and peptides because of its ease of use, low cost, and high complexity.^17,18^ Phage display screen leverages the unique characteristic of bacteriophages, bacteria-infecting viruses, to selectively amplify antigen-specific antibodies by displaying foreign amino acids on their surface while packaging the corresponding coding gene. In addition, bacteriophage replicate to large number and in large quantities.^14^ Despite its pioneering status and widespread use, phage display faces limitations in mammalian systems. While invaluable for in vitro screening, in vivo screens for cell and tissue targeting remain challenging due to unusual clearance mechanisms and concerns regarding immunogenicity and toxicity.^19–21^ The complexity of mammalian tissues (with their structural variations, diverse post-translational modifications, and intricate molecular interactions) presents challenges that conventional display technologies cannot readily overcome.^17^

EVs are a heterogeneous population of lipid-bound nano-size particles secreted by all cell types that offer distinct advantages over conventional library screens and position them for development as a superior screening tool.^22–24^ They mediate intercellular communication by transferring genetic materials, lipids, and proteins, with diverse roles documented in immune regulation^25^, antigen presentation^26^, tissue repair and regeneration^27^, cancer^28^, and neurodegenerative diseases^29^. This role in communication suggests a number of functional readouts that could be built into EV display screens where effective delivery of specific types of cargo results in a detectable signal. EVs demonstrate unique kinetics, biodistribution patterns, functionality, and clearance pathways that synthetic lipids and nanocarriers cannot replicate.^4,30^ Most importantly, they avoid the toxicity and immunogenicity associated with lipid-based nanocarriers and viral delivery systems.^2,31^ While some EVs naturally exhibit tissue tropism, controlling and enhancing this targeting specificity has remained challenging. By adding affinity molecules (such as antibody mimetics or peptides) to the EV surface, cellular affinity and targeted delivery have been altered both in cultured cells and in vivo.^5,8,32–35^.^32^

Despite significant advances in EV research, a systematic approach for screening and selecting molecules that can direct EVs to specific cellular targets in vivo has not been described, hindering the development of precisely targeted therapeutic delivery systems. Building on our previous work, we established the technological foundation for such a strategy that is possible due to two key innovations: (1) the development of SLiCE technology for creating recombinant DNA libraries in mammalian cells with exceptional efficiency^36^, and (2) the engineering of EVs displaying monobodies with nanomolar affinity through lactadherin fusion constructs, demonstrating enhanced cellular targeting capabilities.^32^

The VesicleVoyager platform is based on engineering EVs to display monobodies (small, single-domain binding proteins) on their surface while packaging the encoding DNA within. This creates a self-contained screening system where the phenotype (surface-displayed monobody) and genotype (encoding DNA) remain physically linked. Through iterative rounds of selection and amplification, we can identify specific monobodies that direct EVs to desired tissues, enabling the enrichment of EV populations carrying specific targeting molecules. Using both fluorescence and bioluminescence readouts of selection with sequence validation, we demonstrated successful in vitro and in vivo screening of these EV-displayed monobodies and identification of unique tissue-targeting monobodies.

To further understand the stochastic relationship between displayed monobodies and their encoding DNA within EVs, we developed a Markov chain model that captures the selection dynamics and predicts enrichment patterns. This mathematical framework provides insights into the efficiency of each selection step and explains how successful targeting molecule enrichment occurs despite the imperfect correlation between phenotype and genotype. Here we detail the development, validation, applications, and mathematical modeling of this novel platform.. This innovative approach addresses a critical need for targeted delivery systems that can navigate the complexities of mammalian biology, and has transformative potential for drug discovery, diagnostics, and personalized medicine applications where precise tissue targeting is essential for therapeutic success.

## Materials and Methods

### Reporter construct

The cloning methods and the backbone constructs were adopted from our previous work.^5^ Synthetic double-stranded DNA coding for NanoLuc,^37^ CD63 (Gene bank accession number: NP_001771), and ThermoLuc^38^ were purchased from (Twist Bioscience) to generate pcS-NanoLuc-C1C2 and pcS-ThermoLuc-CD63.^8,32^ Briefly, we amplified the backbone sequence from pcS-mCherry-C1C2 (Addgene #178425) and inserted the codon humanized NanoLuc, CDD63, and ThermoLuc fragments.

### Library DNA preparation

The methods and the backbone constructs were adopted from our previous work.^8,32^ All primer sequences used in this study are listed in Table S1. Briefly, the library backbone was amplified from template pcS-RDG-C1C2 (Addgene #200163) using primers Lib-BB-F and Lib-BB-R, followed by the restriction digest by DpnI, to eliminate template DNA. Monobody library fragments coding for variable loop regions from Hydrophilic Fibronectin Library second generation^39^ containing 44 bp at the 3’-end and 45 bp overlap at the 5’-end were amplified using primers Lib-IN-F and Lib-IN-R^39^ and fused to the backbone using <u>S</u>eamless cloning <u>Li</u>gation <u>C</u>ell <u>E</u>xtract (SLiCE) method.^8,32^ Following the cleanup using QIAquick PCR purification kit (QIAGEN) and the concentration measurement by the Qubit dsDNA BR Kit (Invitrogen), the assembled DNA was electroporated into electrocompetent *E. coli* cells (NEB) and pre-cultured at 37°C for 1 hour without antibiotics, then cultured in the Lauria Broth (LB) containing 100 μg/mL ampicillin for 8 hours at 37°C in a shaking incubator. The library DNA was extracted using the Midiprep kit (QIAGEN), and concentration was determined by NanoDrop (Thermo Fisher Scientific).

### Cell Culture and Treatment

Cell lines were obtained from American Type Culture Collection (ATCC) and routinely tested for mycoplasma contamination. Lines included: HEK293T (Human Embryonic Kidney cell line), A431 (Human carcinoma cell line), and MCF-7 (human breast cancer cell line). Cells were cultured in high-glucose DMEM (Gibco) supplemented with 100U/mL penicillin, 100 µg/mL streptomycin, and 10% (v/v) fetal bovine serum (FBS, Gibco), and maintained in a humidified incubator with 5% CO_2_ at 37°C.

For EV production, HEK293T cells were seeded at 1.5×10^6^ in 10 cm tissue culture dishes 24 hours prior to transfection. For transfection, 10 μg DNA was mixed with in-house PEI (polyethylenimine) reagent prepared from PEI (Sigma 408727) at a DNA:PEI ratio of 1:2.5 (μg:μL) in non-supplemented DMEM, pulse-vortexed for 30 seconds, incubated at room temperature for 10 minutes and added to the cells.^8^ After 24 hours of incubation with 5% CO_2_ at 37°C, cells were rinsed once with PBS, and culture media was replaced with 20 mL of DMEM supplemented with Insulin-Transferrin-Selenium (ITS) (Corning), 100 U/mL penicillin and 100 μg/mL streptomycin (conditioned media) and incubated for another 24 hours for library EV generation. EVs were co-labeled with imaging molecules by co-transfecting 5 μg of monobody-display plasmid and 5 μg of imaging plasmid (pcS-NanoLuc-C1C2, pcS-mCherry-C1C2, pcS-ThermoLuc-CD63) per dish.

### EV isolation

EVs were purified from conditioned media by differential centrifugation. Briefly, culture media was centrifuged at 600g for 30 minutes to remove cells and cellular debris, and the supernatant was further centrifuged at 2000g for 30 minutes to remove apoptotic bodies. The supernatant was then ultracentrifuged in PET Thin-Walled ultracentrifuge tubes (Thermo Scientific 75000471) at 12,000g with a Sorvall WX+ Ultracentrifuge equipped with an AH-629 rotor (k factor = 242.0) for 90 minutes at 4°C to pellet the EVs. The pellet containing EVs was resuspended in 100 µL EV storage buffer.^40^ EVs were also isolated by Tangential Flow Filtration. The supernatant was filtered by MICROKROS 20CM 0.65UM MPES (Repligen, C06-E65U-07-S), and concentrated by MICROKROS 20CM 0.05UM PS (Repligen, C02-S05U-05-S).^9^ Concentrated EVs were rinsed with EV storage buffer and collected in 1mL of EV storage buffer.^40^

### DNase I Treatment of EVs

5 µL of Monobody Library EVs were incubated at room temperature for 15 minutes with 1 U of DNase I (Zymo Research) and DNA Digestion Buffer. The plasmid DNA was then isolated from the EVs using Qiamp Miniprep kits and quantified by qPCR.

### Quantitative Real-time Polymerase Chain Reaction (qPCR)

qPCR was performed using Dream Taq DNA polymerase (ThermoFisher). Each reaction contained 200 µM dNTPs, 500 nM each of forward/reverse primer, 400 nM probe (Table S1), 0.5 U DreamTaq DNA polymerase, 1x Dream Taq buffer A and 1 µL sample DNA in a total reaction volume of 10 µL using CFX96 Touch Real-Time PCR Detection System (BIO-RAD). The PCR amplification cycle was as follows: 95°C for 2 min; 40 cycles of 95°C for 20 seconds, 65°C for 30 seconds. The pDNA copy number were determined by absolute quantification using the standard curve method, and the copy number of EV-encapsulated pDNA per vesicles was calculated based on nanoparticle tracking analysis (NTA) and qPCR results.

### Western Blotting

EVs were denatured at 70°C for 10 minutes in 1x NuPAGE LDS Sample Buffer (Thermo Fisher Scientific), separated on 4–20% Mini-PROTEAN® TGX™ Precast Protein Gels (BioRad), and transferred to Polyvinylidene fluoride or polyvinylidene difluoride (PVDF) membranes using CAPS-based transfer buffer. Membranes were blocked with EveryBlot Blocking Buffer (BioRad) for 2 hours and then incubated with primary antibodies at 4°C overnight. Following three washes with TBS containing 0.1% Tween 20 (TBST), membranes were incubated with horseradish peroxidase-conjugated secondary antibodies for 2 hours at room temperature. After three additional TBST washes, protein bands were visualized using SuperSignal West Pico PLUS chemiluminescent substrate (Thermo Scientific) and imaged with ChemiDoc Imaging System (BioRad). Primary antibodies used: anti-HA (Sigma Aldrich, H3663), anti-CD63 (Thermo Fisher, 10628D), anti-ALIX (Proteintech, 12422-1-AP), TSG101 (Abcam, ab125011), Calnexin (Abcam, ab133615), and GAPDH (Cell Signaling Technology, #2118). Secondary antibodies used: Goat anti-Mouse IgG (H+L) Highly Cross-Adsorbed Secondary Antibody, HRP (Invitrogen, A16078) and Goat anti-Rabbit IgG (H+L) Highly Cross-Adsorbed Secondary Antibody, HRP (Cell Signaling Technology, A16110).

### In vitro EV library screening

For in vitro screening, A431 cells were seeded at 0.3×10⁶ cells/well in 6-well plates 24 hours prior to EV treatment. Cells were treated with 2.0×10⁷ library EVs in 2 mL media for 30 minutes at 37°C. Following PBS washing to remove residual EVs, cells were harvested using trypsin. Plasmid DNA was isolated using a modified protocol for plasmid isolation from organ homogenates using QIAprep Spin Miniprep Kit,^41^ and used as template for the next round of library DNA preparation. This screening process was repeated for 5 rounds to enrich targeting monobody sequences.

### In vitro Bioluminescence assay

A431 cells were seeded at 0.02×10^6^ cells/well in 96-well plates (UV-Star® Microplate, 96 well, COC, F-Bottom (Chimney Well), uClear®, Clear; Greiner Bio-one) 24 hour prior to EV treatment. For the assay, 5×10^6^ of NanoLuc co-labeled EVs (generated by co-transfection with EV-displayed NanoLuc constructs) were added to wells in triplicate. After incubation at 37°C, cells were washed twice with PBS to remove unbound EVs. 50 µL of 1 μg/mL Coelenterazine-H (CTZ; Regis Technologies) was added to each well immediately before imaging. Luminescence was recorded using an in vivo imaging system (IVIS; Spectrum Perkin Elmer) and the particle numbers emitting equal amounts of luminescence/radiance (photons/sec/cm^2^/sr) were calculated.

### Confocal Microscopy

Co-labeled engineered EVs (eEVs) were prepared by co-transfection as described above. A total of 1 × 10^5^ each of A431 and MCF-7 cells were mixed and seeded in 8-well chamber slide (0030742036, Eppendorf, Germany) 24 hours before EV treatment. These co-cultured cells were incubated with 2.0 × 10^7^ monobody-mCherry co-labeled eEVs for 10 min, followed by three PBS washes to remove unbound EVs. Cells were fixed with 4% PFA at room temperature for 15 minutes, washed with PBS containing 0.1% Tween20 three times, and blocked with blocking buffer (1% BSA in PBS) for 60 min. Cells were then incubated with primary antibody (CST 4267T, Cell Signaling Technology, Danvers, MA, USA) in a humidified chamber for 60 minutes at room temperature. After three 5-minutes PBS washes, cells were incubated with secondary antibody (CST 4412, Cell Signaling Technology, Danvers, MA, USA) for 1 hour at room temperature in the dark. After three additional PBS washes, cells were incubated with DAPI for 10 minutes at room temperature. Slide were mounted with mounting medium (00-4958-02 Fisher Scientific, USA) and fluorescence images were taken at 60× magnification using confocal laser scanning microscopy (A1 HD25/A1R HD25 confocal microscope, Nikon, Japan).

### Super Resolution Microscopy

Isolated monobody Library EVs were analyzed with EV Profiler V2 Kit for Nanoimager (ONI) following manufacture’s protocol. Imaging data was analyzed by CODI software (ONI).

### Next-Generation Sequencing

The sample for next-generation sequencing (Illumina MiSeq) were prepared by PCR amplification using a primer pair (CS1-LibHA-F, CS2-G4S-R) with sequencing indices. PCR products were quantified using Qubit before sequencing. All samples were normalized to the same concentration, and agarose gel electrophoresis confirmed product size. Sequencing was performed at the MSU Genomics Core facility using MiSeq Reagent Kit v3 for 250 bp paired-end reads. The generated FASTQ files were extracted, processed, and clustered by sequence similarity using custom software^42^.

### In vivo EV library screening

In this study, 8- to 12-week-old female Balb/cJ mice from Jackson Laboratories were used for animal experiments. Animals were housed in the University Laboratory Animal Resources Facility, and all procedures were performed with approval from the Institutional Animal Care and Use Committee of Michigan State University. Approximately 1× 10^9^ monobody library EVs in EV storage buffer^40^ were injected into mice intravenously (IV). One hour after administration, mice were sacrificed, and visceral organs (heart, lung, liver, kidney, pancreas and spleen) were excised and homogenized using Bulk Ceramic Beads 2.8mm (Fisher Scientific) and BeadBug 6 Microtube Homogenizer (Benchmark Scientific). Plasmid DNA was isolated from organ homogenates using QIAprep Spin Miniprep Kit using modified protocols for plasmid isolation from mammalian cells^8,41^, and used as template for the next round of library preparation. This screening process was repeated for 5 rounds to enrich targeting monobody sequences. The enriched variants were re-cloned into the EV display construct for individual monobody characterization.

### In vivo and ex vivo imaging

To quantify relative detection levels in *ex vivo* studies, bioluminescence per 1x 10^8^ particles was measured for each EV sample. Approximately 1× 10^9^ monobody EVs co-labeled with NanoLuc were injected intravenous into Balb/c mice (n=3). One hour after administration, mice received an intraperitoneal injection of CTZ (10 µg/g), 10 minutes prior to bioluminescence imaging (BLI) using an In Vivo Imaging System (IVIS; Revvity, previously PerkinElmer). Mice were anesthetized with isoflurane and imaged intact. Following in vivo imaging, the mice were sacrificed and visceral organs (heart, lungs, liver, kidneys, pancreas, and spleen) were dissected and imaged using an IVIS. Bioluminescence in each organ was quantified, and the amount of detection was normalized as the relative detection value to the initially introduced bioluminescent material. For in vivo imaging with ThermoLuc, approximately 2× 10^10^ Monobody EVs co-labeled with ThermoLuc were injected intravenous into Balb/c mice (n=3).

### Markov chain modeling of the selection process

To analyze the stochastic relationship between EV surface-displayed monobodies and their packaged plasmid DNA, we developed a discrete-state Markov chain model of the selection process.^43^ The model consists of seven states representing key stages in the selection cycle: (1) Initial Library, (2) EV Production, (3) DNA Packaging, (4) Target Binding, (5) DNA Recovery, (6) Enriched Library, and (7) Selection Loss. Transition probabilities between states were estimated from experimental data as follows:

The probability of successful monobody display on EVs (*p_display_*) was calculated based on super-resolution microscopy analysis, which showed that approximately 80% of EVs displayed monobodies. The DNA packaging efficiency (*p_packaging_*) was derived from DNase-resistant plasmid DNA quantification, with an average of 1.5 DNA copies per EV, resulting in a packaging probability of approximately 0.6. Binding probabilities (*p_binding_*) were estimated from time-course experiments with A431 cells, with target-binding monobodies showing approximately 0.3 probability of specific binding. DNA recovery efficiency (*p_recovery_*) following tissue binding was calculated at 0.5 based on qPCR quantification of DNA recovered from target organs.

The overall single-round selection efficiency was calculated as:

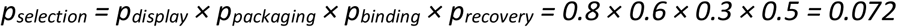

For stochastic linkage analysis between surface-displayed monobodies and their encoding DNA, we calculated the probability of correct linkage (*p_linkage_*) as:

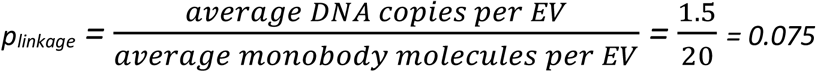

The probability that an EV contains any plasmid DNA (*p_anyDNA_*) was calculated using a Poisson distribution with λ = 1.5:

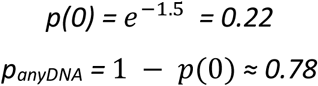

For enrichment simulations, we used a computational model that incorporated both deterministic selection based on transition probabilities and stochastic processes reflecting the imperfect linkage between phenotype and genotype. The model was implemented in Python, and all simulations were performed for 5 consecutive rounds to match our experimental protocol.

### Simulation of monobody variant enrichment

To model the enrichment of specific monobody variants over successive selection rounds, we employed a competitive binding model that incorporates both selection pressure and initial variant frequency.

For a given monobody variant with initial frequency *f_0_*in the library, the frequency *f_i_* after round i was calculated as:

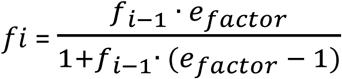

where *e_factor_* is the enrichment factor calculated as:

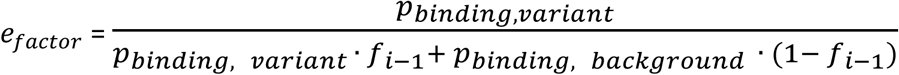

Here, *p_binding, variant_* is the binding probability of the variant of interest (e.g., 0.6 for E626), and *p_binding, background_* is the average binding probability of the remaining library (e.g., 0.1 for non-specific binders).

This model allows simulation of both rapid enrichment of high-affinity variants and competitive exclusion of non-binders. For organ-specific targeting, we extended the model to simulate biodistribution using experimental data. For example, the pancreas-enriched Pan-P5 library exhibited a ∼4-fold increase in pancreatic accumulation compared to the initial library (P0), which was incorporated into the simulation to project shifts in organ-level distribution of EVs.

## Results

### EV-based monobody display screening strategy and design

Our EV-monobody screening platform design builds on the method of decorating EV surfaces using EV-surface display constructs.^32,44^ We used a yeast monobody library^39^ to generate the DNA for an EV-display monobody library with a diversity of 4.2×10^9^. Negative depletion magnetic bead sorting was done against streptavidin-coated beads removed non-selective binders (approximately 1×10^7^ to 1×10^8^ monobodies). DNA from the remaining 10^9^ monobody variants was cloned into the EV display vector using SLiCE, transfected into HEK293 cells, and EVs were recovered as the displayed monobody library. After isolating the EV-monobody library from conditioned media, we quantified vesicle numbers and assessed plasmid DNA loading prior to screening in cultured cells or animals.

For repeated selection, cells or organs were harvested, DNA extracted, monobody-coding regions amplified by PCR, and re-cloned into the EV-display backbone. The process was repeated 5 times to ensure co-selection of enriched monobodies and their encoding DNA (Fig. 1).

**Figure 1.**
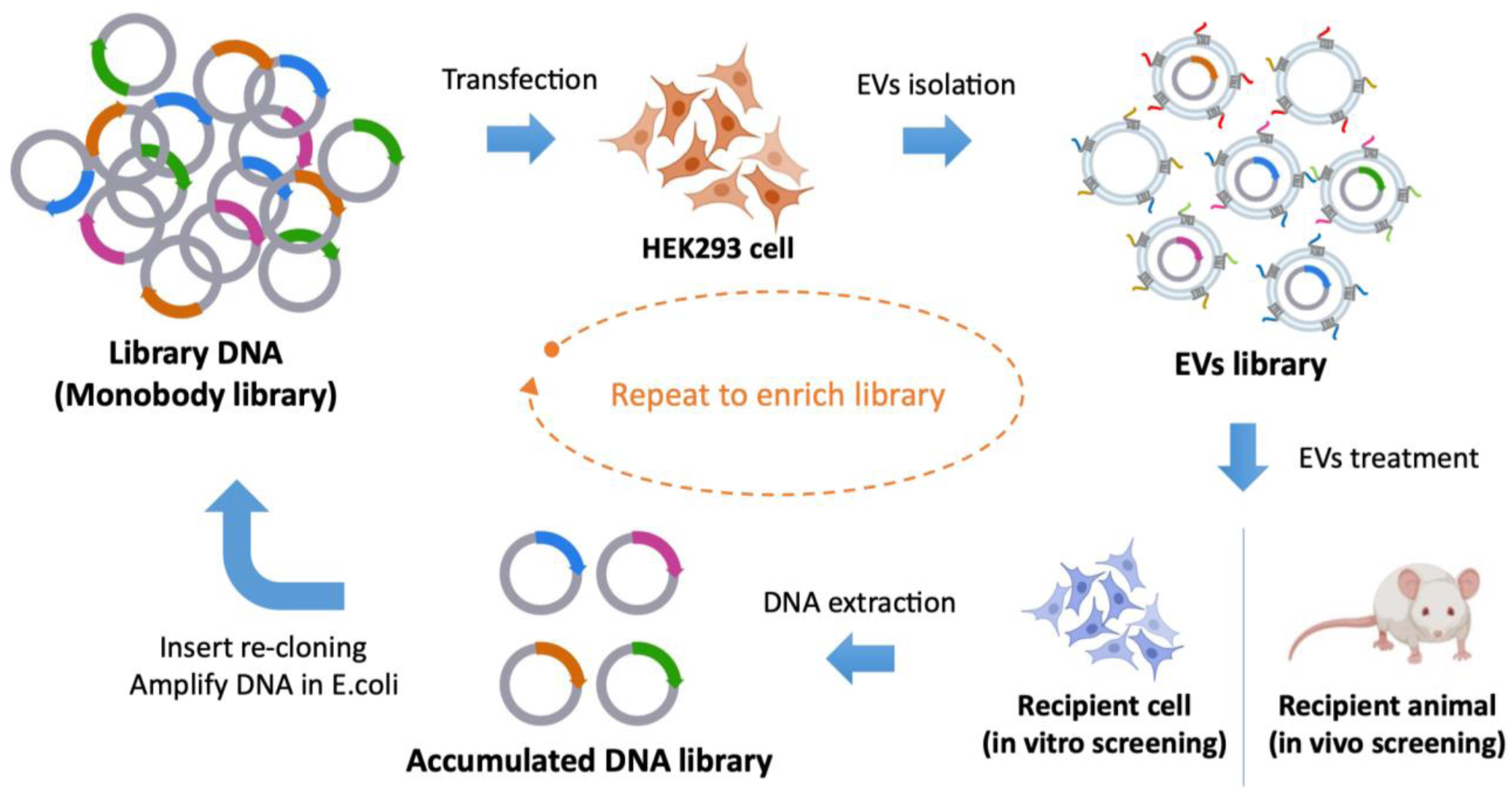
Schematic illustration of monobody-EV library screening strategy. Overview of EV-based monobody screening processes, involving monobody library pDNA generation, pDNA transfection to EV donor cells (HEK293T), EV isolation from the cell culture media, EV treatment or administration, DNA extraction from cell or organ, monobody amplification and re-cloning to generate enriched monobody library pool. This image was created with BioRender.com.

### Monobody Library EVs from HEK293T cells display monobody proteins while packaging and safeguarding pDNA

Large and small EVs can be separated using differential centrifugation.^45^ Engineered EVs (eEVs) contain the pDNA used for transfection.^8^ To identify an EV population with higher pDNA content, we isolated large EVs (lEVs) at 12,000 x g and small EVs (sEVs) at 100,000 x g for 90 minutes from the supernatant after centrifugation of lEVs.^8^ Most pDNA was present in the lEVs separated at 12,000 x g (Fig. S1), confirming previous reports.^7^ We therefore focused on lEVs for our study.

For quality control, each EV library batch was characterized using nanoparticle tracking analysis (NTA) and quantitative PCR (qPCR). NTA consistently revealed peak EV sizes of 120-140 nm (Fig. 2A,B). To confirm that pDNA was protected by EVs, we treated samples with DNase I, which degrades free-floating DNA to undetectable levels, then quantified pDNA.^8^ Figure 2C illustrates the distribution of monobody-coding pDNA per EV, with an average packaging efficiency of approximately 1.5 copies per particle. Western blot analysis demonstrated enrichment of EV markers (CD63, TSG101, and Alix), successful monobody display (HA), and absence of the cell-specific marker calnexin (Fig. 2D).^46^ Super-resolution microscopy confirmed monobody protein display through co-localization with EV markers (CD63 and CD9) and HA-tag (Fig. 2E-H), with approximately 20 monobody molecules expressed per EV particle (Fig. 2I).

**Figure 2.**
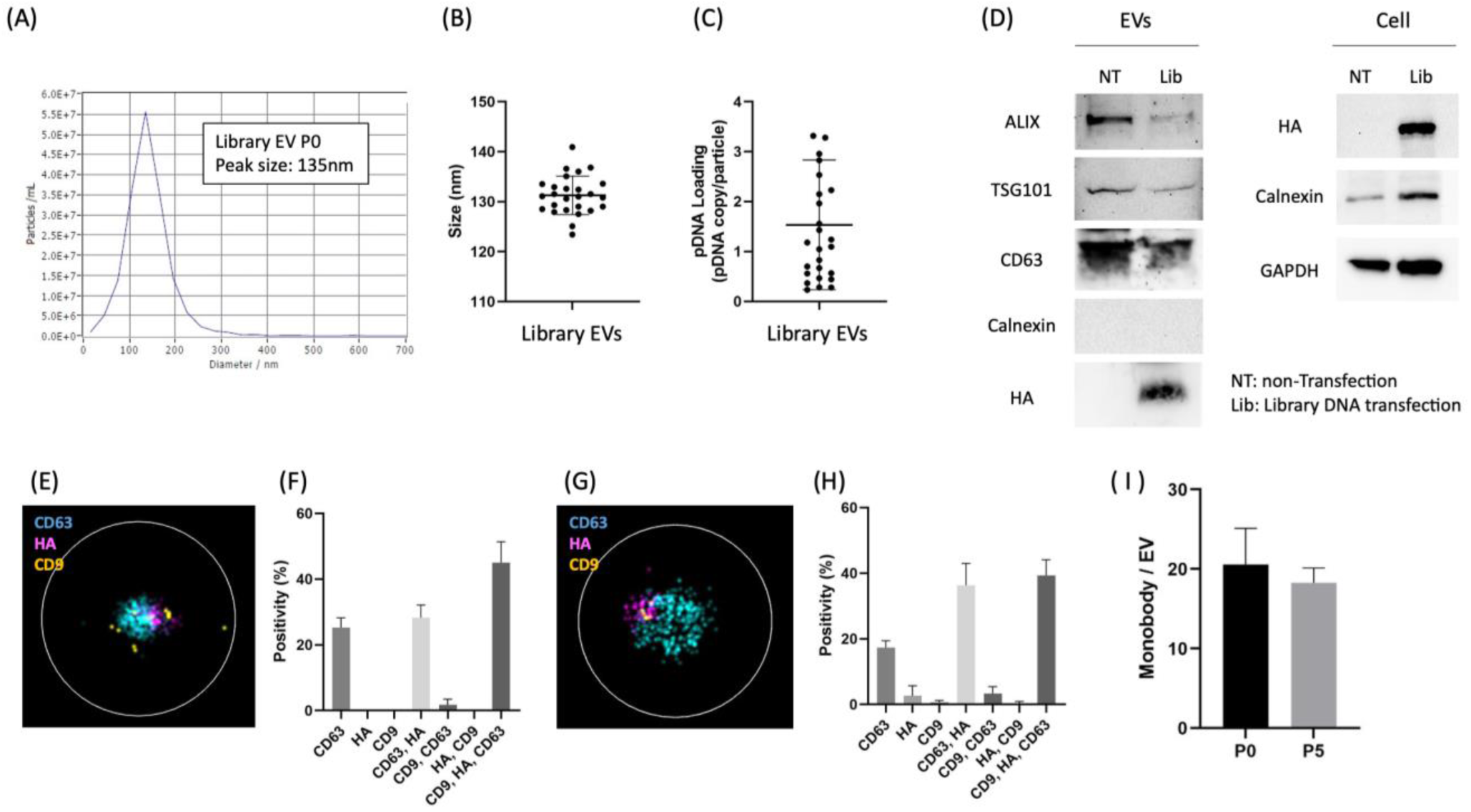
EV characterization showing successful EV isolation, pDNA loading, and surface display of monobody proteins. (A) Size distribution, (B) Peak size distribution of multiple samples, (C) Amount of pDNA loading in eEV, (D) immunoblot analysis, (E) Representative image of eEV before screening and (F) population analysis, and (G-H) after five rounds of screening, respectively. (I) Number of monobody molecules expressed on the EV surface.

While we could not establish a direct one-to-one link between specific surface-displayed proteins and their corresponding packaged plasmid DNA within individual EVs, our iterative selection process demonstrated that targeting monobodies were successfully enriched through multiple rounds of screening. This suggests that despite the stochastic nature of the initial library, the selection process effectively identified and amplified functional targeting molecules.

### In Vitro Enrichment of Cell Type-Specific Monobody Sequences

We conducted time-course experiments using A431 EGFR+ cells treated with an EGFR-specific monobody (E626)^47^, non-binder control (RDG), and co-labeled EVs to determine the optimal treatment duration for in vitro screening (Fig. S2). We observed time-dependent increases in pDNA uptake with both monobodies, but EVs labeled with anti-EGFR monobody E626 exhibited significantly higher pDNA uptake compared to non-binder RDG or co-labeled EVs. This difference was evident at 15 minutes and more pronounced at 30 minutes.

Based on these results, we initially selected a 30-minute treatment duration as optimal. However, since 15-minute treatments also showed significant differences, we compared both timepoints directly. To assess enrichment of high-binder monobody sequences, we spiked 1% each of E626 (positive control) and RDG (negative control) into the monobody library. After three screening rounds, E626 was enriched at both treatment durations, but more efficiently with 30-minute treatments (Fig. S3). The complete list of sequence data is available on GitHub https://github.com/HaradaLabMSU/EV-Library.

We performed five rounds of screening with 30-minute EV treatments. qPCR analysis demonstrated that E626 was significantly enriched in rounds one through three and stabilized in rounds four and five (Fig. 3A). The occurrence frequency of each unique sequence from NGS analysis was tabulated for each experimental replicate and panning round (p1-p5). The ratio between E626 and RDG sequences is plotted in the figure 3B. An increase in the E626/RDG ratio—indicating enrichment of E626 and depletion of RDG—occurred in five out of six independent screening experiments by the third round. The decrease in unique sequences and increase in prevalent-to-rare sequence ratios suggest library collapse during iterative panning (Fig. S4). This library collapse is expected in successful screening, where high-performing sequences should appear with increasing frequency (Fig. 3B).

**Figure 3.**
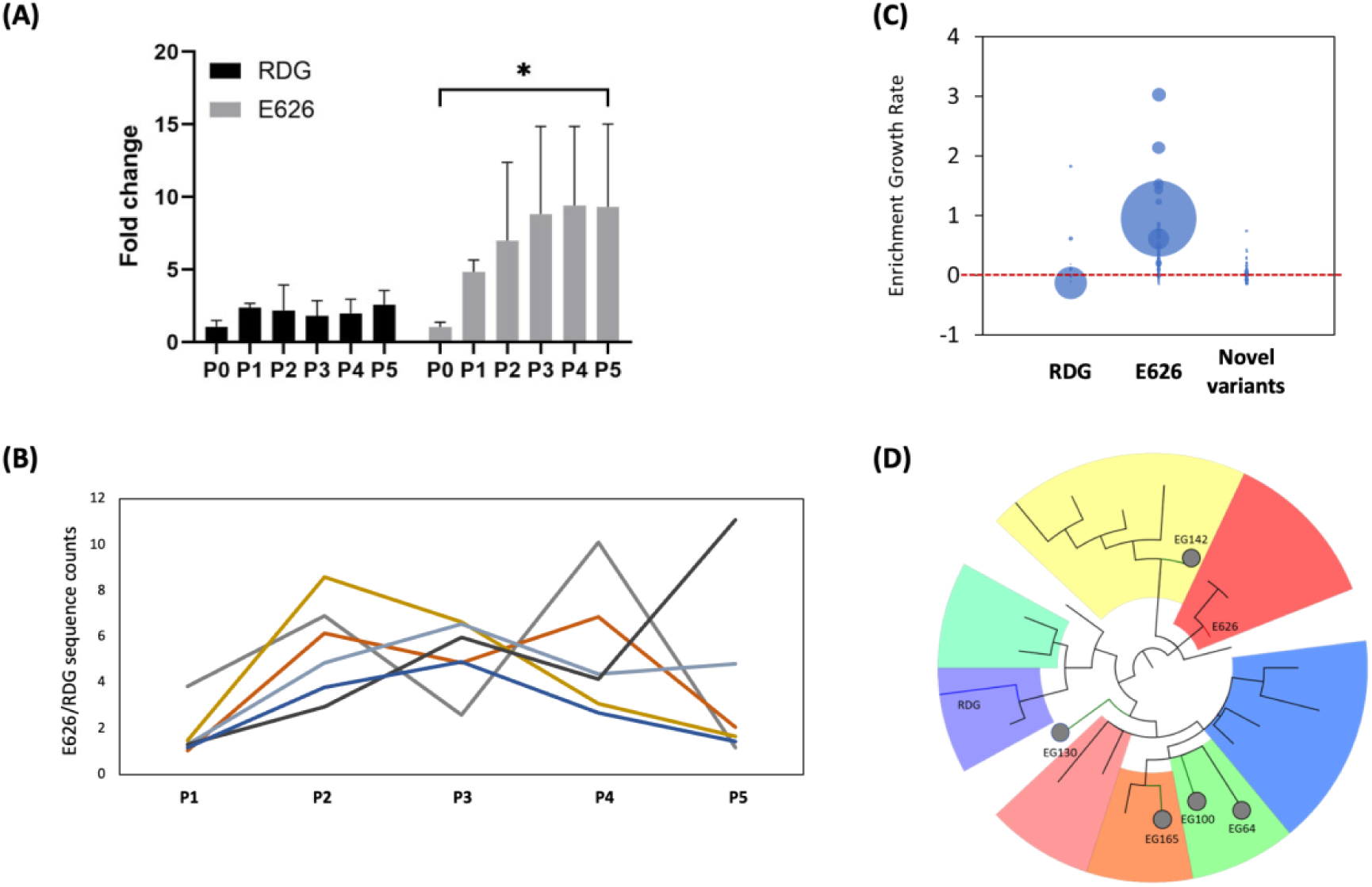
Enrichment of target sequences on in vitro EV-Monobody library screening. **(A)** Validation of fold change of RDG and E626 in A431 cells by qPCR. **(B)** The ratio of E626 (positive control) to RDG (negative control) sequences quantified for each of five passages across six separate campaigns. **(C)** Enrichment growth rate of each variant calculated using: Final count = initial count × Exp (enrichment growth rate × time). **(D)** Phylogenetic tree showing relatedness between positive control (E626), negative control (RDG), and novel lead monobody variants EG64, EG100, EG130, EG142, and EG165.

Furthermore, novel variants were enriched from a naïve monobody library. In Fig. 3C, each dot represents a unique monobody variant, with dot size proportional to sequence frequency across all replicates. The phylogenetic tree (Fig. 3D) shows relatedness between positive control (E626), negative control (RDG), and novel lead monobody variants (named EG64, EG100, EG130, EG142, and EG165). NGS revealed 562 unique protein variants, which were clustered based on amino acid sequence similarity using CD-Hit (90% threshold). Sequences from the resulting 25 clusters were used to generate the phylogenetic tree. We identified E626 variants (with mutations likely introduced during PCR or cellular replication) and novel clones, suggesting competitive selection of high-affinity monobodies. A panel of variants showing the highest enrichment over multiple passages was selected for further characterization.

To better understand the dynamics of the selection process, we developed a Markov chain model that captures the stochastic relationship between displayed monobodies and their encoding plasmid DNA. Our model predicts that despite the imperfect correlation between phenotype (displayed proteins) and genotype (encoding DNA), the competitive advantage of high-affinity binders during multiple rounds of selection enables successful enrichment of target-binding sequences. The model accurately predicted the enrichment trajectory of E626, closely matching our experimental observations of increasing E626/RDG ratios over five rounds of selection.

### Enhanced Monobody Sequences Exhibit Strong Binding Affinity to Target Cells

We investigated the binding properties of enriched sequences to target cells by re-cloning five high-binder candidates (EG64, EG100, EG130, EG142, and EG165) into an EV-display backbone (Table S2). A bioluminescence-based binding assay using EVs displaying monobodies and NanoLuc revealed that two monobodies (EG130 and EG142) and the positive control (E626) exhibited significantly higher binding relative to controls and other binders (Fig. 4A).

**Figure 4.**
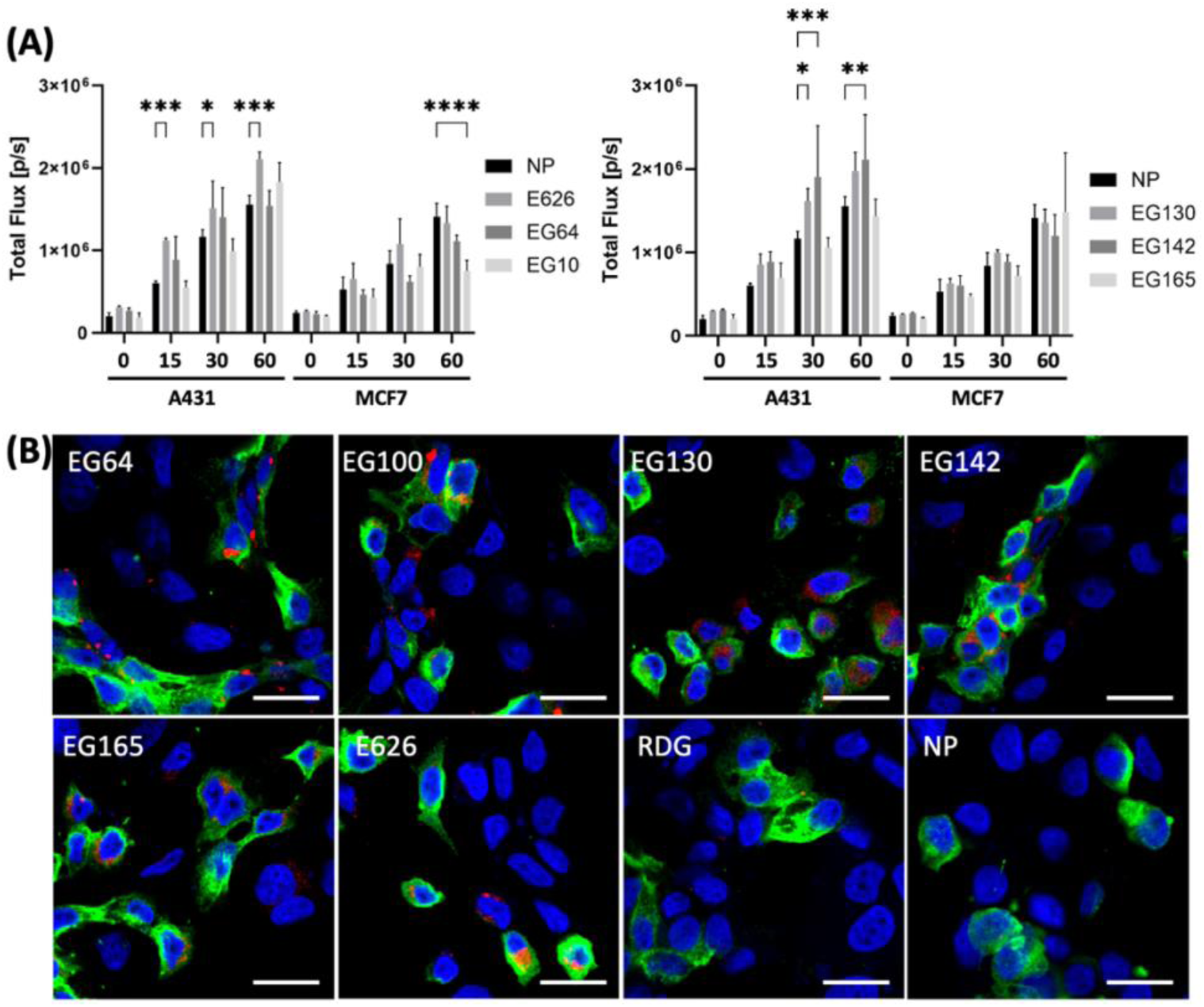
Evaluation of selected clones using bioluminescence imaging (BLI). **(A)** A431 and MCF7 cells were treated with monobody-displayed EVs co-labeled with NanoLuc. Total photon flux (p/s) from EVs bound to cells was quantified using IVIS. Values represent mean ± SD (n=3). Two-way ANOVA was used to assess time course effects. Significance against 0 min is expressed as: * p ≤ 0.05, ** p ≤ 0.01, *** p ≤ 0.001, and **** p ≤ 0.0001. NP: non-peptide, non-binder negative control. **(B)** A431 and MCF-7 cells were co-cultured and treated with Monobody-mCherry co-labeled EVs for 10 min. Binding was assessed by confocal laser scanning imaging of EVs (red), anti-EGFR antibody (green), and DAPI nuclear staining (blue). Scale bar: 25 μm.

To evaluate specific binding capability at the single-cell level, we used a co-culture system for confocal microscopy. We generated mCherry-coated EVs co-labeled with monobodies and treated co-cultured A431 and MCF7 cells for 10 minutes, followed by anti-EGFR antibody and DAPI staining. E626 and high-binder candidates selected by in vitro screening demonstrated specific binding to A431 cells (Fig. 4B, Fig. S5). No specific binding was observed with RDG or NP (negative controls). Notably, EG64, EG100, and EG165, despite not showing statistical superiority in the bioluminescent assay, bound to EGFR-positive cells at high frequency and EGFR-negative cells at low frequency. These findings align with the binding assay results and confirm selection of biomolecules specific to recipient cells through in vitro EV library screening.

### In Vivo Screening of EV-Based Monobody Libraries

To determine optimal circulation time for in vivo screening, we administered library EVs to mice via tail vein and euthanized animals at different timepoints (1, 4, and 24 hours). We isolated pDNA from various organs (liver, kidney, pancreas, and spleen) to determine which duration produced the highest pDNA yield. Recovery varied by organ and timepoint, with average recovery decreasing from 1 to 4 or 24 hours across all organs (Fig. S6A). Based on these findings, we used a 1-hour circulation time for subsequent library screening.

We confirmed successful pDNA recovery and monobody fragment amplification from these organs (Fig. S6B). Previous studies demonstrated that pDNA introduced via eEVs can be detected in heart and lungs.^8^ Therefore, we selected six key organs (heart, liver, lung, pancreas, kidney, and spleen) to test for specific EV accumulation.

The monobody EV library was injected into mice via tail vein and allowed to circulate for 1 hour. After sacrifice, target organs were excised, pDNA extracted from each organ, and organ-enriched monobody sequences amplified and re-cloned into the EV-display backbone. Individual EV libraries were prepared from each organ, then pooled for subsequent screening rounds (Fig. S7). Three independent rounds of in vivo screening were conducted, and library DNA from the fourth and fifth rounds underwent next-generation sequencing, revealing several enriched monobody sequences within each organ’s DNA library (Table S3).

### Pancreas-Enriched Monobody Library EVs Exhibit Targeted Accumulation

The pancreas is a challenging organ to target therapeutically, making it an important focus in this research. To validate in vivo enrichment of pancreas-targeting monobodies, we compared biodistribution of the initial monobody library EVs (P0) with pancreas-enriched monobody library EVs (Pan-P5). The Pan-P5 library consisted of pooled monobody-library pDNA after five independent in vivo screening rounds.

Both P0 and Pan-P5 monobody library EVs were co-labeled with NanoLuc and administered intravenously. Sequential imaging over 30 minutes revealed similar initial biodistribution patterns (Fig. S8). However, at the 1-hour timepoint, ex vivo imaging demonstrated significantly increased Pan-P5 monobody library EV accumulation within the pancreas compared to P0, providing robust evidence of pancreas-specific monobody enrichment through in vivo screening (Fig. 5A).

**Figure 5.**
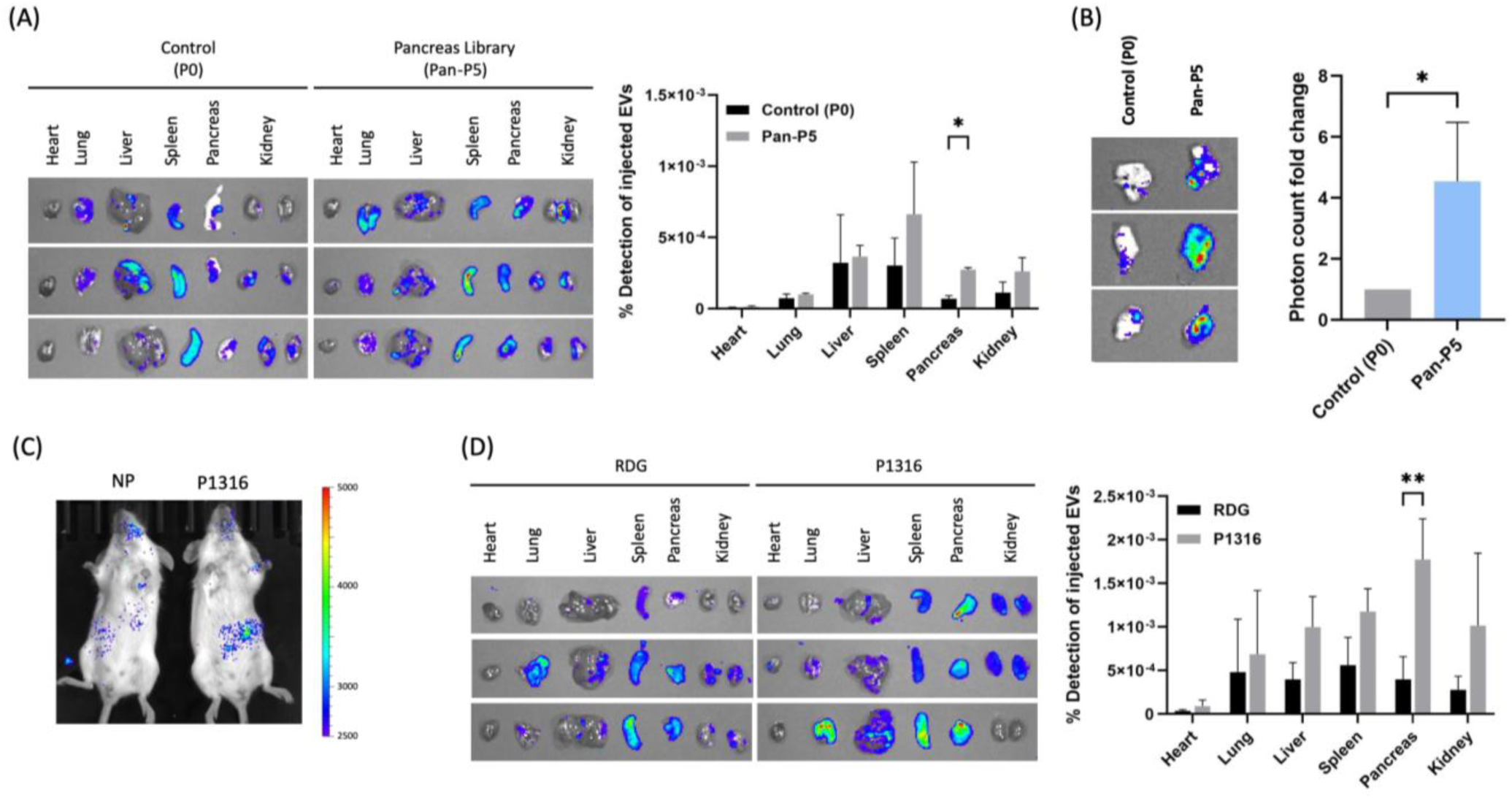
Five rounds of in vivo screening created a library with high pancreatic accumulation. EVs were co-labeled with NanoLuc and each Library DNA (P0, Pan-P5), high binder candidate P1316 or non-binder RDG. Equal numbers of particles were introduced into mice by IV injection, and ex vivo imaging was performed after 1 hour of circulation. **(A)** Ex vivo images of each organ, comparison of percentage of detected BLI from injected EVs, **(B)** ex vivo image of pancreases, accumulation degree of pancreatic library relative to control (original library). **(C)** Biodistribution of P1316-EVs co-labeled with ThermoLuc-CD63. **(D)** Ex vivo image of each organ, comparison of percentage of detected BLI from injected EVs. Unpaired t-test was used to evaluate the enrichment in each organ. Significance is expressed as follows: * p ≤ 0.05, ** p ≤ 0.01.

To quantify this enrichment, we normalized signals from each organ to the input. The Pan-P5 library EV signals were approximately twice as high as those of P0. In a direct pancreas-to-pancreas comparison, the Pan-P5 signal was approximately four times higher than that of the P0 library (Fig. 5B).

Next, we evaluated the ability of individual monobody sequences (whose enrichment was confirmed by sequence analysis) to accumulate in the pancreas. Four high-binder candidates were cloned into an EV-display construct and preliminarily assessed for pancreatic binding capacity. These high-binder candidate EVs, co-labeled with NanoLuc protein by co-transfecting EV-displayed Nanoluc pDNA with the EV-displayed monobody pDNA into producer cells. The high-binder, Nanoluc EVs were introduced into mice via tail vein injection, and organ accumulation was evaluated through ex vivo imaging. Among the candidates, P1316 demonstrated particularly high pancreatic accumulation (Fig. S9).

We further evaluated P1316 using EVs co-expressing the monobody and a ThermoLuc-CD63 fusion protein, whose expression is specific for intracellular delivery. P1316-EVs showed a signal vicinity to the pancreas region compared to the control, NP (Fig. 5C, Fig. S10). Furthermore, ex vivo evaluation using NanoLuc confirmed significant accumulation in the pancreas (Fig. 5D), supporting the successful enrichment of organ-specific biomolecules through our in vivo screening approach.

### Markov chain modeling reveals selection dynamics of EV-displayed monobodies

To further understand the stochastic relationship between surface-displayed monobodies and their packaged DNA, we developed a Markov chain model of the selection process (Fig. 6A). This model enabled us to quantify the efficiency of each step in the selection cycle and predict enrichment rates for specific monobody variants.

**Figure 6.**
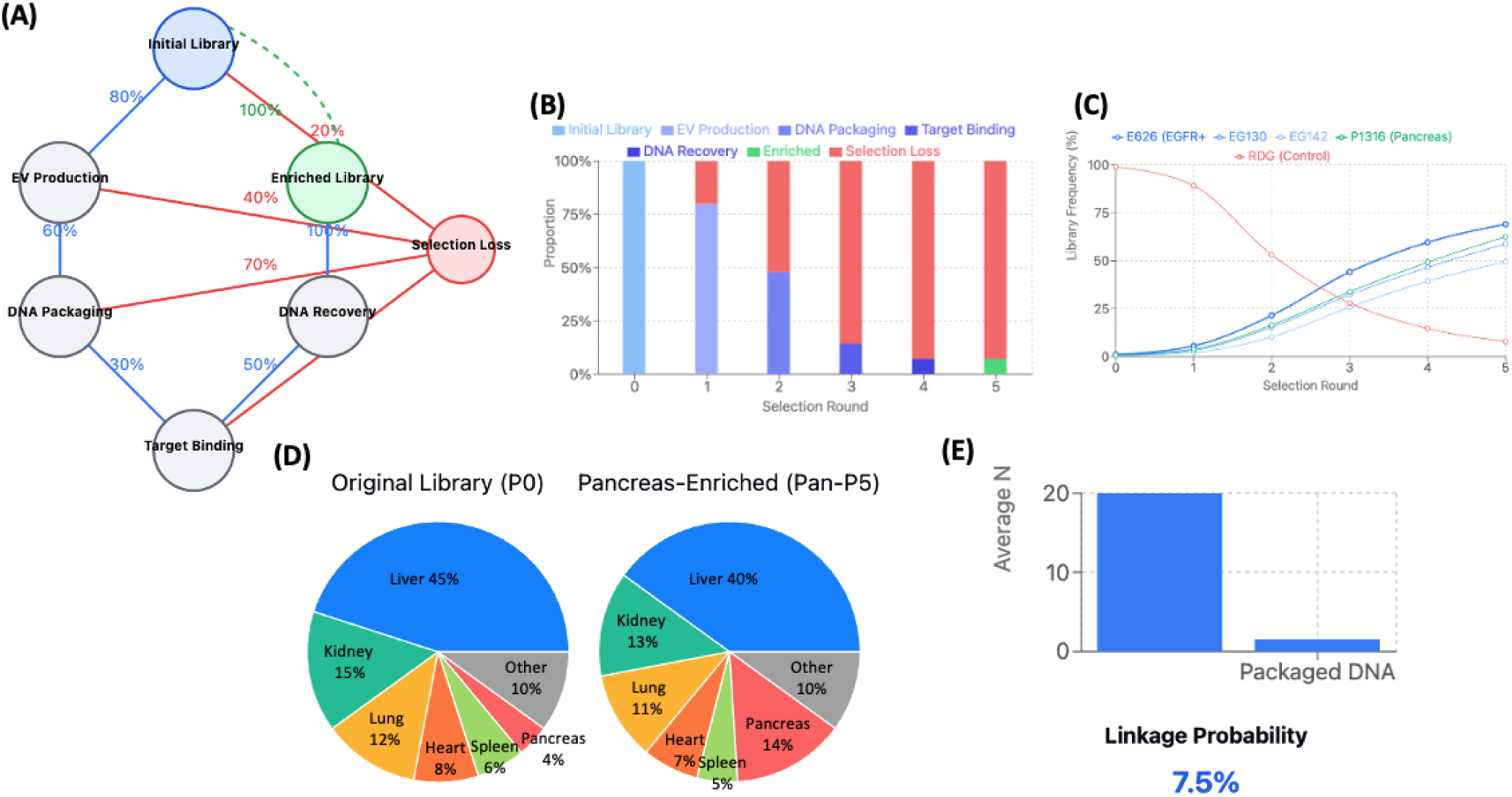
Markov chain model of EV-displayed monobody selection process. **(A)** Schematic representation of the Markov chain model, showing the seven states and transition probabilities between states. Each state represents a key stage in the selection process, with arrows indicating possible transitions. **(B)** Simulated population distribution across states after each selection round, showing the progressive enrichment of the library and cumulative loss through multiple stages. **(C)** Enrichment of E626 monobody frequency over five rounds of selection from an initial 1% to 69%, demonstrating the effectiveness of the selection process despite imperfect genotype-phenotype linkage. **(D)** Comparison of organ distribution profiles between the initial library (P0) and pancreas-enriched library (Pan-P5), showing the shift in targeting specificity following selection. Error bars represent standard deviation from three independent simulations. **(E)** Stochastic relationship between displayed monobodies and packaged DNA. Each EV displays approximately 20 monobody molecules but contains an average of only 1.5 copies of plasmid DNA, resulting in a ∼7.5% probability of correct linkage between a displayed monobody and its encoding DNA. Despite this imperfect correlation, the iterative selection process successfully enriches target-binding sequences. Error bars represent standard deviation from three independent simulations.

The model revealed that approximately 7.2% of the initial library successfully completes a single round of selection, with the remaining 92.8% lost at various stages (Fig. 6B). The greatest loss occurs during the target binding step (70% loss), followed by DNA packaging (40% loss), DNA recovery (50% loss), and display failure (20% loss). Despite these sequential losses, the model showed that target-binding monobodies can be successfully enriched through multiple rounds of selection.

Our stochastic linkage analysis showed that while each EV displays approximately 20 monobody molecules on its surface, it contains on average only 1.5 copies of plasmid DNA. This results in a 7.5% probability of correct linkage between a displayed monobody and its encoding DNA. Despite this imperfect correlation, our simulations demonstrated that the competitive advantage of high-affinity binders during the selection process can overcome the stochastic nature of genotype-phenotype linkage.

Simulation of E626 enrichment (initially present at 1% of the library) showed progressive enrichment to 5.5%, 21.4%, 44.1%, 59.6%, and 69.0% through five rounds of selection (Fig. 6C). This corresponds to a change in the E626:RDG ratio from 0.01 to 2.23, closely matching our experimental observations. The model thus confirms that despite the stochastic nature of monobody display and DNA packaging, our iterative selection process effectively enriches target-binding sequences.

For organ-specific targeting, our model predicted that a 4-fold increase in pancreatic accumulation (as observed for Pan-P5 compared to P0) would shift the organ distribution profile, increasing pancreatic targeting from 4% to 14% while proportionally decreasing targeting to other organs (Fig. 6D). These predictions correlated well with our ex vivo imaging results for P1316, validating the model’s utility in predicting organotropic targeting.

## Discussion

In the development of new drugs and targeted therapies, relevant screening techniques for selecting tissue and cell targeting molecules from a combinatorial library will enable directed delivery and reduce off-target toxicity. Screens that inherently use particles in the selection process should select for targeting proteins that can direct nanoparticles and other large cargos to target cells and tissues. While phage display^48^, yeast display libraries^49^, and various screening methods have successfully demonstrated biomolecule selection^20^, our EV-display library screens extend peptide and protein screening to in vivo assays with greater relevance to targeted drug and nanoparticle delivery.

The key challenge in our approach was establishing the correlation between the displayed surface proteins and the packaged DNA within individual EVs. Our Markov chain model quantified this relationship, revealing that each EV displays approximately 20 monobody molecules but contains an average of only 1.5 copies of plasmid DNA, resulting in a 7.5% probability of correct linkage between a displayed monobody and its encoding DNA in the original EV library, with greater probabilities expected in sequentially selected pools as the library complexity collapses. Library convergence to fewer unique sequences increases the frequency of identical sequences, where higher frequencies indicate more effective selection. The distribution of these sequence frequencies provides quantitative data for modeling selection dynamics and assessing screening efficiency. Although we could not demonstrate a direct one-to-one relationship between surface-displayed monobodies and their corresponding encoding DNA in single EVs, the iterative selection process effectively enriched targeting monobodies. This suggests that despite the initially stochastic nature of protein display and DNA packaging, the selection pressure applied through multiple rounds of screening successfully identifies functional targeting molecules.

Screening display libraries in relevant mammalian ecosystems using naturally occurring bionanoparticles (EVs) reveals targeting peptides that direct particles, and perhaps small molecules, to target tissues. The EV structure, a lipid bilayer membrane encapsulating various biomolecules, allows cells to be used as packaging factories to produce engineered particles with defined cargo and specific targeting molecules on their surface.^2^ In this way, the natural function of EVs for intercellular communication can be adapted for use as delivery vehicles. Our approach takes advantage of modified cargo (pDNA) to correlate with modified EV surfaces, enabling effective in vivo screening. Our Markov model demonstrates that this stochastic approach is robust, with each round of selection providing approximately 7.2% successful recovery of the library while enriching target-binding sequences.

Various methods for modifying the EV surface have been reported.^6^ In our previous studies, we demonstrated targeting by EV surface display using the C1C2 domain of human lactadherin.^8,32^ Our engineered EVs (eEVs) shield at least one transfected pDNA molecule per particle from external environmental factors. Additionally, they express an average of 20 monobody molecules on their surface. Using these characteristics, we designed an EV library screening methodology similar to phage display screens but adapted to the stochastic relationship between DNA cargo and surface markers.

To our knowledge, this is the first report of EVs being used in a biomolecule screening platform, either in culture or in vivo. As an initial demonstration, we performed in vitro screening using A431 cells. Both qPCR and NGS evaluations confirmed enrichment of the anti-EGFR monobody E626, which was spiked in as a positive control. Our study involved five rounds of screening, but E626 enrichment was confirmed after just the first round, including in our preliminary experiments. This suggests that library collapse occurs quickly in vitro, and selection may be possible with fewer rounds, making the screening process more cost-effective and efficient than anticipated.

The improved efficiency may relate to the EV type selected. Characterization of the library EVs showed an average particle size of approximately 130 nm containing, on average, 1.5 molecules of pDNA per particle. The detection of typical EV markers (CD63, ALIX, and TSG101) along with the engineered HA marker confirmed that the collected particles were EVs, with 80% displaying monobodies on their surface. While there are few reports quantitatively evaluating EV labeling efficiency, our results may serve as a guide for future research.^50–52^ Our single EV imaging studies confirmed that monobodies can be displayed on the EV surface in multiple copies, which may account for our better-than-expected results and provide a significant advantage over phage display, where typically only 5 copies of the same peptide are present on each particle.^20^

The process by which pDNA is packaged into eEVs is likely due to cytoplasmic DNA not being tolerated by cells, with EV packaging serving as one mechanism to remove this DNA.^53^ The relationship between the expression of EV-targeted proteins (the lactadherin fusion proteins), biogenesis of eEVs, and packaging of the pDNA cargo is likely stochastic. The stochastic relationship we observed between surface-displayed proteins and packaged plasmid DNA aligns with recent findings by Fordjour et al. (2022), who demonstrated that extracellular vesicle cargo proteins can vary by approximately 100-fold from one vesicle to another through a shared stochastic mechanism.^54^ In Markov state models, the system transitions between discrete states with specific probabilities, without requiring a deterministic one-to-one correlation between surface proteins and internal cargo. While each individual EV may contain variable combinations of surface proteins and internal DNA, our repeated rounds of selection effectively collapsed this probabilistic distribution toward EVs containing both the targeting monobody on the surface and its encoding DNA inside. This progressive enrichment can be conceptualized as a directed evolution of the Markov process, where selection pressure drives the system toward specific high-probability states that represent functional targeting EVs.

While we could not establish a direct correlation between displayed proteins and packaged DNA within individual EVs, our results demonstrate that through iterative rounds of selection, we successfully enriched targeting monobodies. The fact that anti-EGFR monobody sequences were enriched in our screening suggests that even without perfect initial correlation between phenotype and genotype in the library, the selection process effectively identifies functional targeting molecules with each round of selection.

Among the five high-binder candidate sequences newly identified by in vitro screening, EG130 and EG142 showed significant affinity to A431 cells in bulk assays. Confocal microscopy confirmed that all sequences were specific to A431 cells, possibly due to the sensitivity of the assay. The pancreatic library that underwent five rounds of in vivo screening showed higher affinity for the pancreas compared to the starting P0 library, and the high-affinity candidates obtained by NGS analysis also demonstrated affinity for the pancreas. This confirms that our in vivo EV library screening successfully selects for tissue-targeting monobodies. However, we observed that even EVs expressing high-affinity monobodies for the pancreas showed some non-specific delivery to other organs, suggesting that further improvements in specificity are necessary for therapeutic applications.

The identification of clinically relevant targeting monobodies presents challenges that can be addressed in several ways. Compared to in vivo screening of phage display libraries, EV library screening avoids potential bystander effects on the mammalian microbiome and reduces risks to the immune system because it utilizes a universal mammalian mechanism rather than a bacterial virus. Once a targeting monobody has been identified, its target protein and epitope can be characterized, and if homologous to human proteins, translation to clinical applications is straightforward.

Alternatively, monobody libraries can be pre-selected against human targets and then screened in vivo to increase the likelihood of finding clinically relevant monobodies. EV libraries could also be screened in humanized mice to identify relevant monobodies. In this study, we used HEK293T cells to produce eEVs, but the intrinsic targeting properties of EVs vary depending on the cell type used for production. Therefore, careful consideration of the cells used as EV factories is important.^55,56^ Furthermore, using disease models as screening hosts could potentially select molecules that directly target disease sites.

## Conclusion

We have developed a screening system that utilizes engineered EVs as a peptide display platform to select molecules that direct nanoparticles to target cells and tissues. If the identified monobodies are used on other nanoparticles, some of the off-target delivery observed in our study would likely be reduced. Alternatively, if synthetic EVs are used, or if natural EVs can be “hardened” to prevent non-specific delivery to non-target sites, monobody-coated nanoparticles could achieve highly selective drug delivery to specific cells and tissues. This technology opens an exciting new approach to drug discovery that combines the advantages of natural EVs with the power of directed molecular evolution. Our mathematical modeling provides a framework for understanding the selection dynamics of EV-displayed monobodies. Despite the stochastic relationship between displayed proteins and packaged DNA, the iterative nature of the selection process enables successful enrichment of targeting molecules. This quantitative understanding of the selection process will guide optimization of future screening campaigns and facilitate the development of more efficient targeted delivery systems.

## Data availability

The datasets generated and/or analyzed in this study are available in the HaradaLabMSU/EV-Library GitHub repository, https://github.com/HaradaLabMSU/EV-Library. GitHub repository is available under the Creative Commons Attribution-NonCommercial-NoDerivatives 4.0 International Public License (“Public License”) at https://github.com/HaradaLabMSU/EV-Library.

## Acknowledgement

We thank Dr. Seock Jin Chung and Mr. Chia-Wei Yang for assisting with animal experiments, the IQ Advanced Molecular Imaging Facility for in vivo and ex vivo imaging, the Center for Advanced Microscopy for confocal imaging, the RTSF Genomics Core for sequencing experiments and Dr. Mark Reimers for statistical analysis. This work was funded in part by 1R21GM154180-01 and 1R01CA286786-01 (MH), and the Michigan Translational Research and Commercialization (MTRAC) program (MH, YKH and CHC), which aims to advance the commercialization of innovative technologies by providing early-stage funding, and the James and Kathleen Cornelius Endowment (CHC, YKH and AVM).

