## supplemental for "VesicleVoyager: In vivo selection of surface displayed proteins that direct extracellular vesicles to tissue-specific targets"

Supplementary Information

Table S1. Primer and Probe sequence

| Name | Sequence (5’-3’) |
| --- | --- |
| Lib-BB-F | GGTGGAGGCGGTTCAGGC |
| Lib-BB-R | agcataatctggaacatcatatggataGGC |
| Lib-IN-F | CCAGCCTCCTCGTCGCCTAT |
| Lib-IN-R | CGATCCGCCACCGCCAGA |
| qAmp-1_F | CTAGAGTAAGTAGTTCGCCAGTTAAT |
| qAmp-1_R | GCTGAATGAAGCCATACCAAAC |
| qAmp-1_P | ATTGCTACAGGCATCGTGGTGTCA |
| qRDG-F | TGTACGCTGTAACCGGTCG |
| qRDG-R | CCTGAGACGGTTTGTCGATTT |
| qRDG-P | TCCAGCCGTCCAATCAGCATCAAT |
| qE626-F | CGGGCCAGGATTATACCATTAC |
| qE626-R | GGTTTATCAATTTCGGTGCGATAG |
| qE626-P | ATGCGGTGACCGATAACAGCCATT |

Table S2. Enriched monobody sequence via in vitro screening

| ID | Amino acid sequence |
| --- | --- |
| EG64 | SSDSPRNLEVTNATPNSLTISWDAPYTHATDGYRITYGETGGNSPSQEFTVPGTTNATISGLKPGQDYTITVYAVSDYDLDSNPISINYRTEIDKPSQ |
| EG100 | SSDSPRNLEVTNATPNSLTISWDAPRYSASGYRITYGETGGNSPSQEFTVPGNNTTATISGLKPGQDYTITVYAVSNHGYSNPISINYRTEIDKPSQ |
| EG130 | SSDSPRNLEVTNATPNSLTISWDAYYYGAYYYRITYGETGGNSPSQEFTVAGSYNTATISGLKPGQDYTITVYAVTNDNVSDSNPISINYRTEIDKPSQ |
| EG142 | SSDSPRNLEVTNATPNSLTISWDNHAYHTCYYRITYGETGGNSPSQEFTVPGYSYATISGLKPGQDYTITVYAVSDVSNDDYESNPISINYRTEIDKPSQ |
| EG165 | SSDSPRNLEVTNATPNSLTISWDAPYIAAGYRITYGETGGNSPSQEFTVPGTTSNATISGLKPGQDYTITVYAVTSYDDNSNPISINYRTEIDKPSQ |

Table S3. Enriched monobody sequence via in vivo screening

| Organ | ID | Amino acid sequence |
| --- | --- | --- |
| Spleen | 1873 | SSDSPRNLEVTNATPNSLTISWDTPANAYYYRITYGETGGNSPSQEFTVPGTYYATISGLKPGQDYTITVYAVTGPNYYSGISNPISINYRTEIDKPSQ |
| Spleen | 2437 | SSDSPRNLEVTNATPNSLTISWDNYDSFRTYYYRITYGETGGNSPSQEFTVPGITNNATISGLKPGQDYTITVYAVSSRNLSNPISINYRTEIDKPSQ |
| Liver | 1046 | SSDSPRNLEVTNATPNSLTISWDYYYCSNYYRITYGETGGNSPSQEFTVPGTNSNATISGLKPGQDYTITVYAVSDANYNWSNPISINYRTEIDKPSQ |
| Liver | 1390 | SSDSPRNLEVTNATPNSLTISWDAPDSTYGYRITYGETGGNSPSQEFTVPRSSYATISGLKPGQDYTITVYAVCSVNPFSNPISINYRTEIDKPSQ |
| Liver | 1739 | SSDSPRNLEVTNATPNSLTISWDAPHQTSGYRITYGETGGNSPSQEFTVPRNTNATISGLKPGQDYTITVYAVGNNNHSNPISINYRTEIDKPSQ |
| Kidney | 2202 | SSDSPRNLEVTNATPNSLTISWDAPCVCTSGYRITYGETGGNSPSQEFTVPGTYNYATISGLKPGQDYTITVYAVSNCNHYNVSNPISINYRTEIDKPSQ |
| Kidney | 732 | SSDSPRNLEVTNATPNSLTISWDAPKSAGGYRITYGETGGNSPSQEFTVPGSNTTATISGLKPGQDYTITVYAVSNCDYSNPISINYRTEIDKPSQ |
| Kidney | 1295 | SSDSPRNLEVTNATPNSLTISWDAPTHAHGYRITYGETGGNSPSQEFTVPRNNTNATISGLKPGQDYTITVYAVSNYNHSHHVSNPISINYRTEIDKPSQ |
| Pancreas | 1316 | SSDSPRNLEVTNATPNSLTISWDDHSTTGYYRITYGETGGNSPSQEFTVPGWISTATISGLKPGQDYTITVYAVPTVGFDSNPISINYRTEIDKPSQ |
| Pancreas | 2002 | SSDSPRNLEVTNATPNSLTISWDAPTYYTYGYRITYGETGGNSPSQEFTVPGYNNSATISGLKPGQDTITVYAVGSDDYYSNPISINYRTEIDKPSQ |
| Pancreas | 1540 | SSDSPRNLEVTNATPNSLTISWDAPDNSNGYRITYGETGGNSPSQEFTVPGSTTATISGLKPGQDYTITVYAVTTQGYFSNPISINYRTEIDKPSQ |
| Pancreas | 1544 | SSDSPRNLEVTNATPNSLTISWDAPDYASGYRITYGETGGNSPSQEFTVPGNNNTATISGLKPGQDYTITVYAVSCYNRSNPISINYRTEIDKPSQ |
| Lung | 1696 | SSDSPRNLEVTNATPNSLTISWDAPAHATGYRITYGETGGNSSSQEFTVPRYNYTATISGLKPGQDYTITVYAVTGYNDNPSNPISINYRTEIDKPSQ |
| Lung | 1904 | SSDSPRNLEVTNVTPNSLTISWDAPAVTVRYYRITYGETGGNSPSQEFTVPGSRSTATISGLKPGQDYTITVYAVTGRDGSPASSRPISINYRTEIDKPSQ |
| Lung | 1842 | SSDSPRNLEVTNATPNSLTISWDNPHISFYYRITYGETGGNSPSQEFTVPRNNYATISGLKPGQDYTITVYAVCDTSPYSNPISINYRTEIDKPSQ |
| Heart | 2332 | SSDSPRNLEVTNATPNSLTISWDAPYSASGYRITYGETGGNSPSQEFTVPGYSNATISGLKPGQDYTITVYAVTAYSHVISNPISINYRTEIDKPSQ |

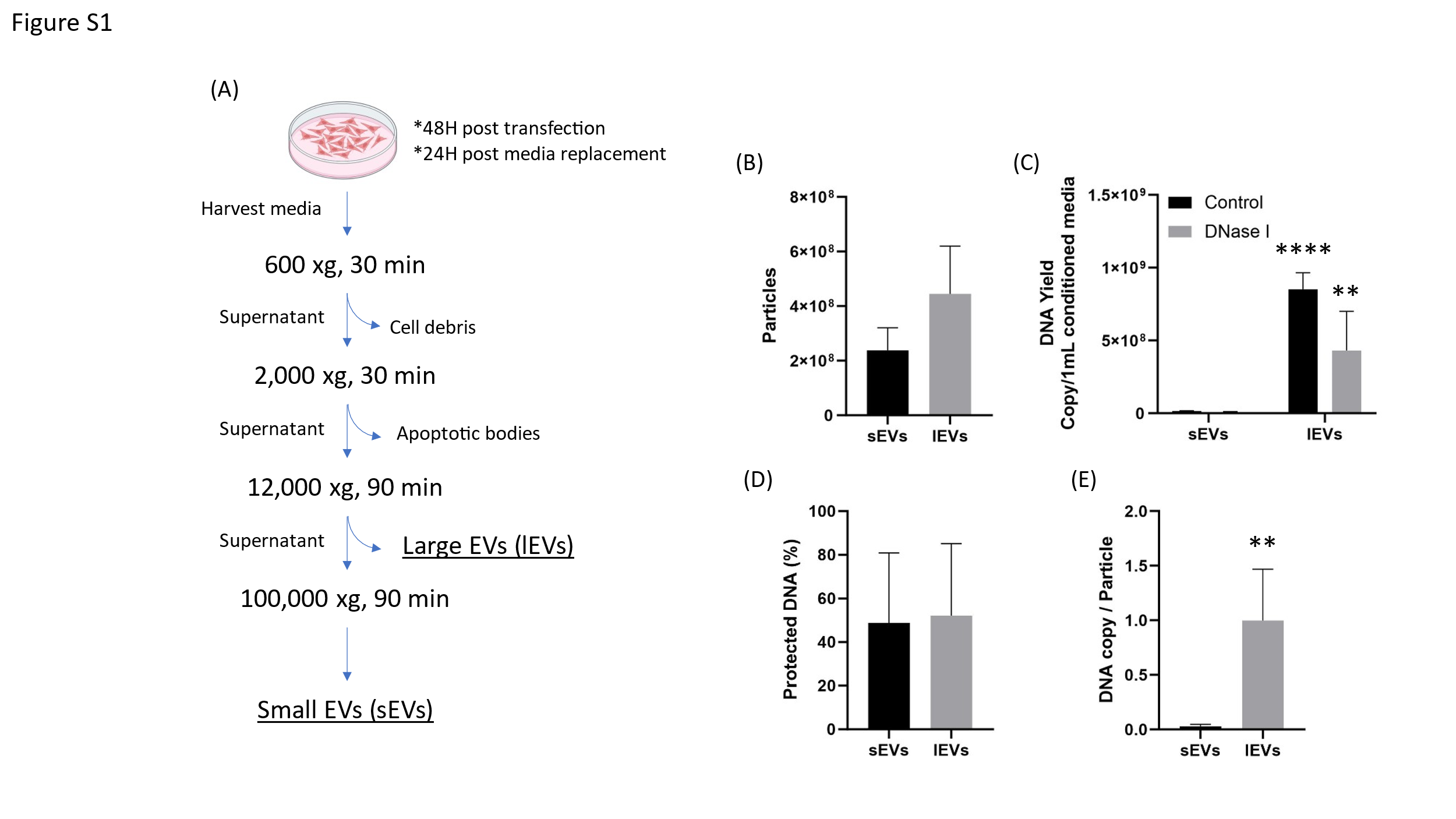

Figure S1. EV isolation process for pDNA richly loading eEVs was established based on differential centrifugation. lEVs and sEVs were separated by different centrifugation speeds and assessed characteristics of (B) particle yield, (C) DNA amount from 1 mL of conditioned media, (D) ratio of encapsulated pDNA inside of eEVs, and (E) pDNA copy number per particle. ** p ≤ 0.01, **** p ≤ 0.0001

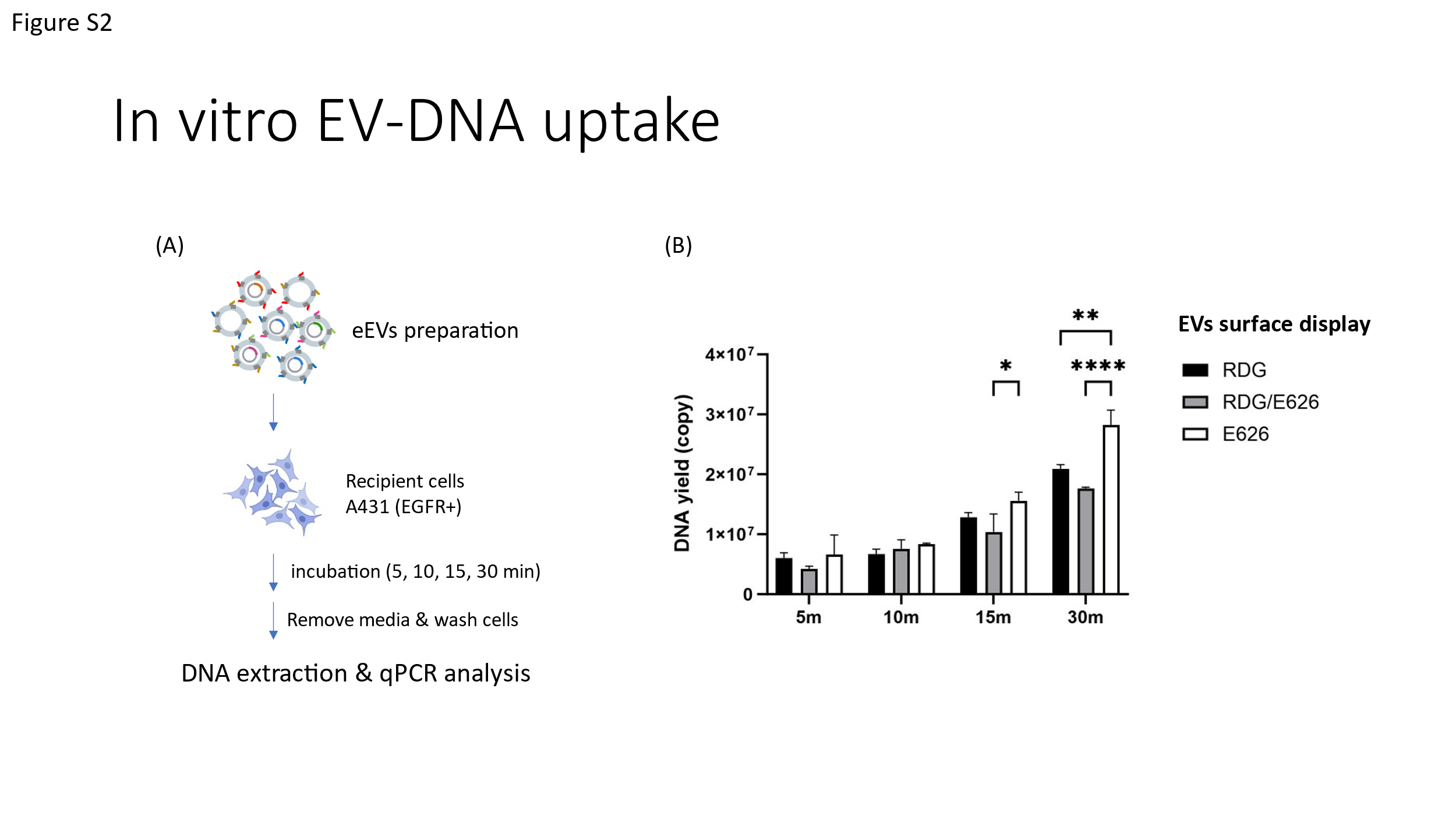

Figure S2. Evaluation of eEV-mediated in vitro uptake of pDNA. A431 cells (EGFR-positive) were treated with eEVs displayed with non-specific Monobody RDG, anti-EGFR Monobody E626, and both on the surface, and uptaken pDNA was quantified. (A) Schematic diagram of the evaluation procedure for in vitro pDNA uptake. (B) Quantification of pDNA taken up into cells. Two-way ANOVA was used to evaluate the effect of the time course in the group. In all figures, significance is expressed as follows: * p ≤ 0.05, ** p ≤ 0.01, and **** p ≤ 0001, if not otherwise specified.

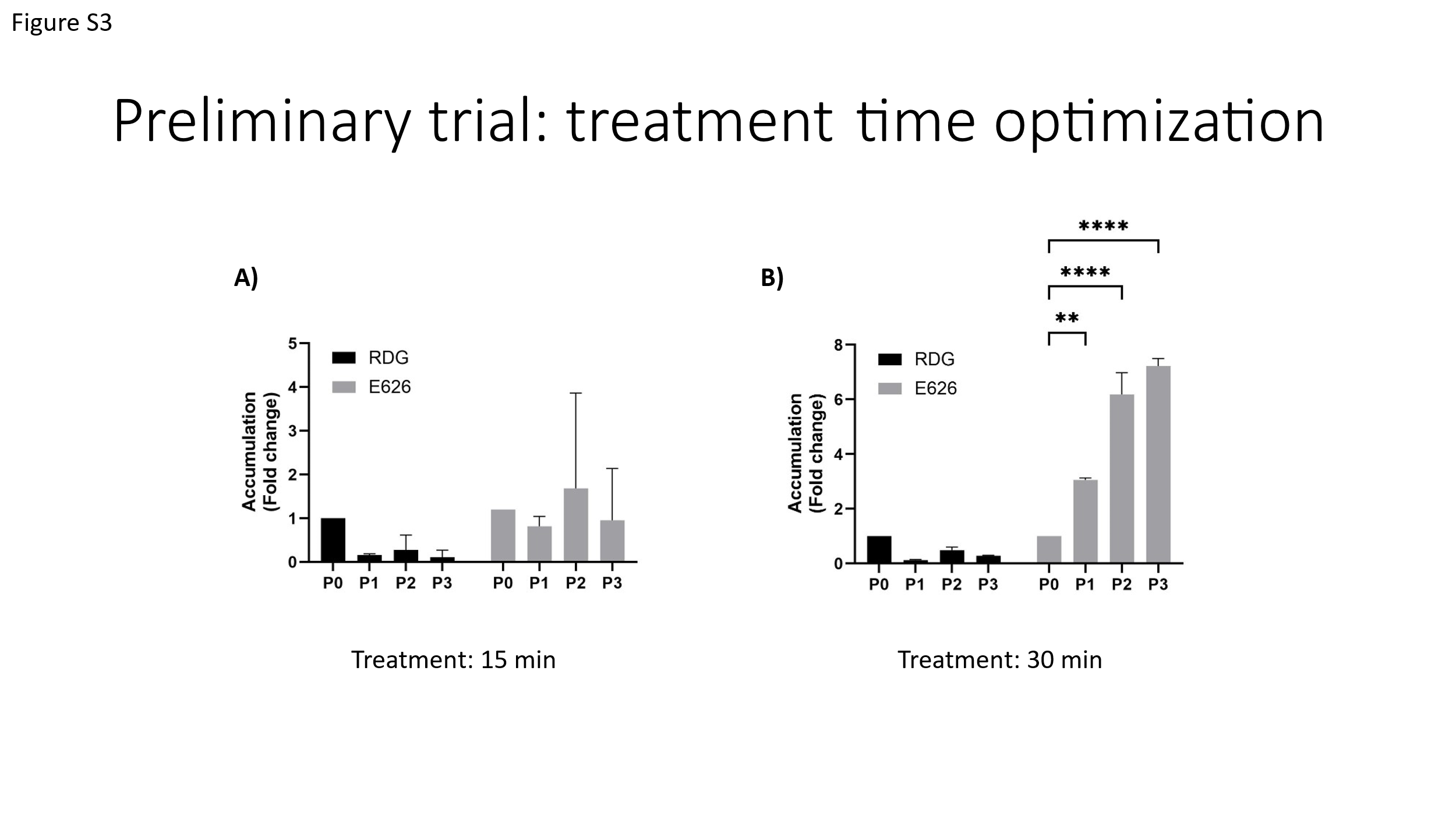

Figure S3. Optimization of EVs treatment time on in vitro EV Library screening. Three rounds of screening were attempted by EVs treatment time (A) 30 min and (B) 15 min. Two-way ANOVA was used to evaluate the effect of the time course in the group. In all figures, significance against P0 is expressed as follows: ** p ≤ 0.01 and **** p ≤ 0001, if not otherwise specified.

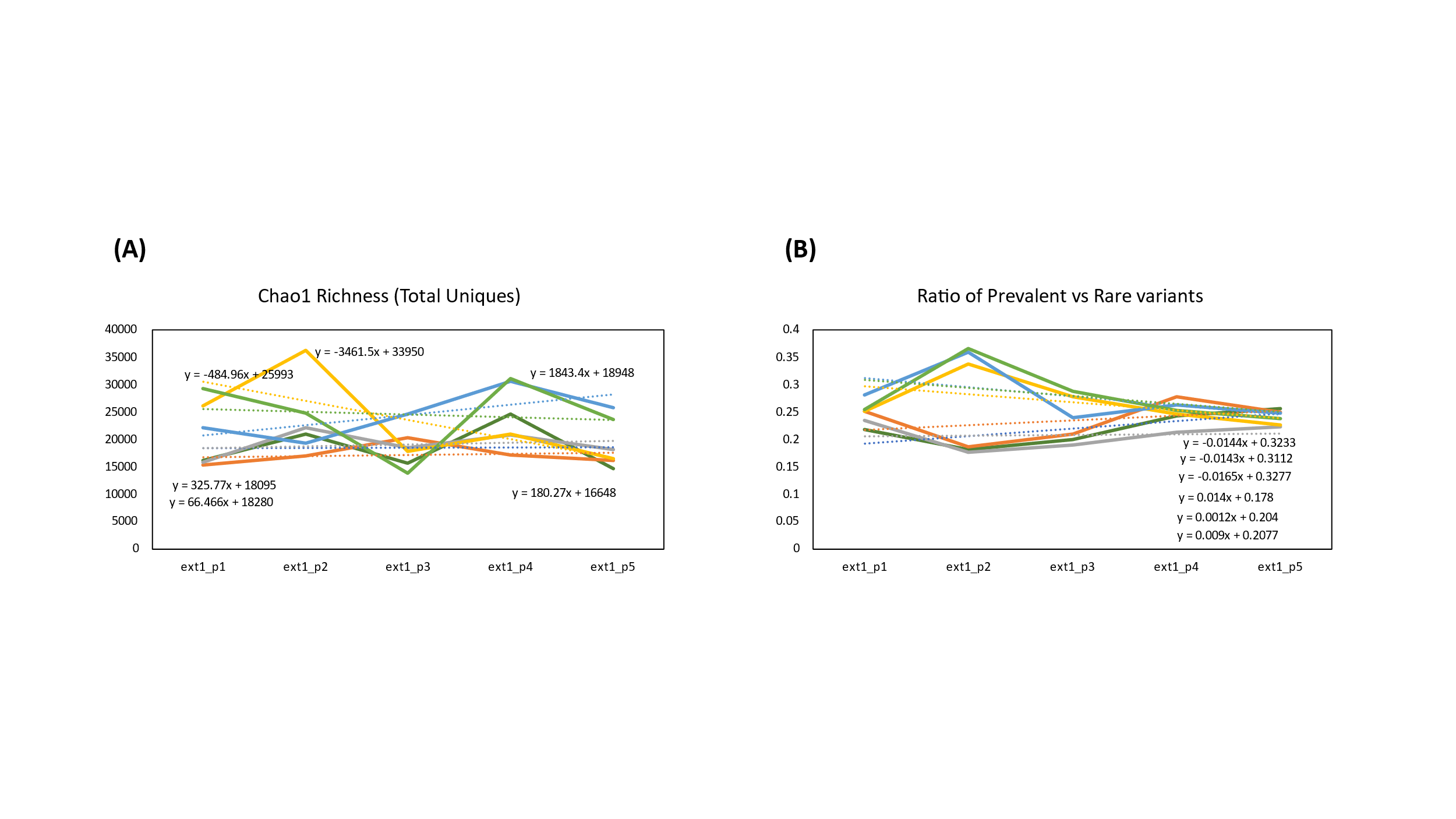

Figure S4. Assessing library complexity by NGS analysis. (A) Chao1 Richness was used to estimate the total number of unique variants. (B) The ratio of prevalent variants and rare variants (using a cutoff of n<3 reads) was calculated for each sample and passage.

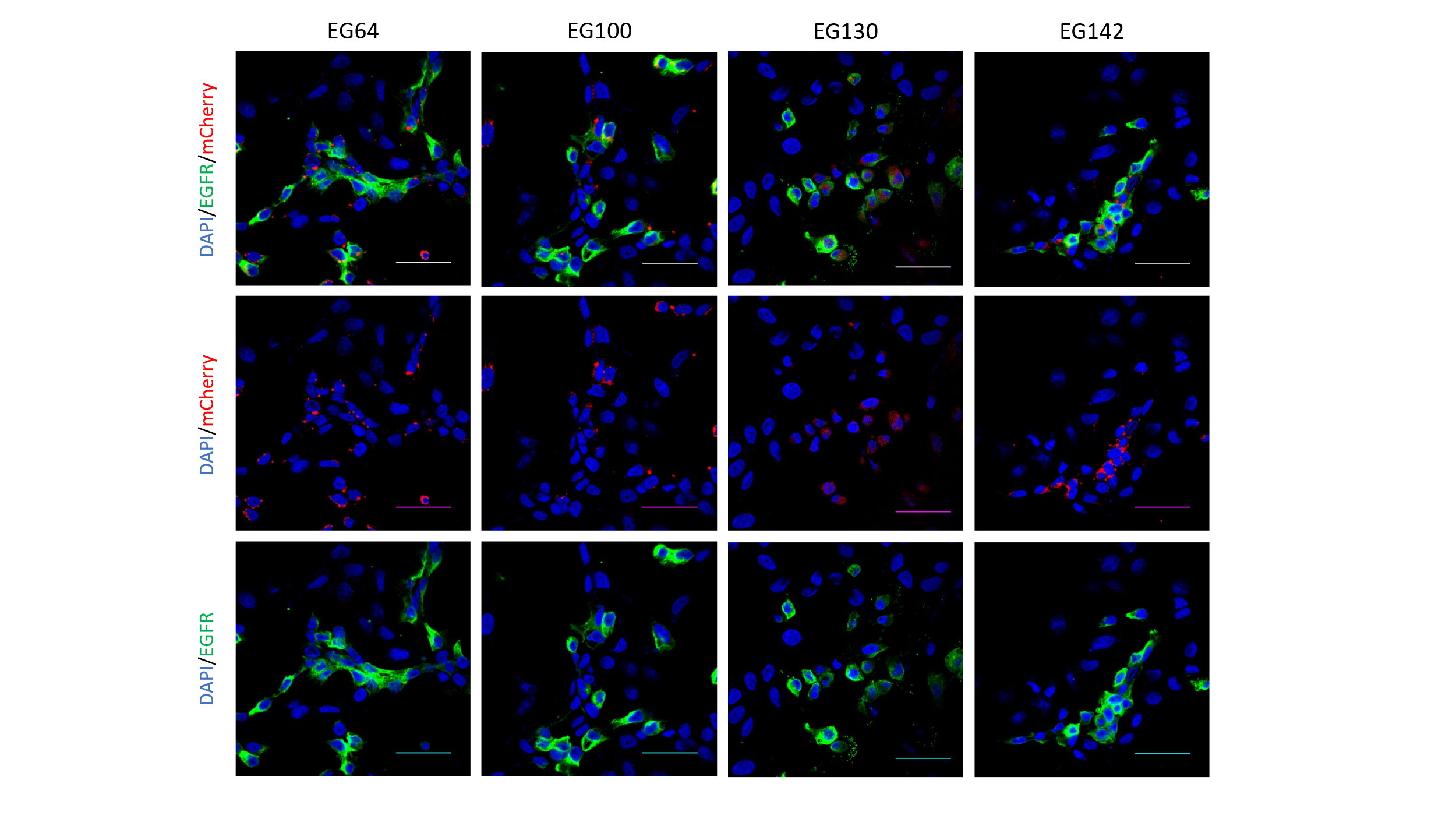

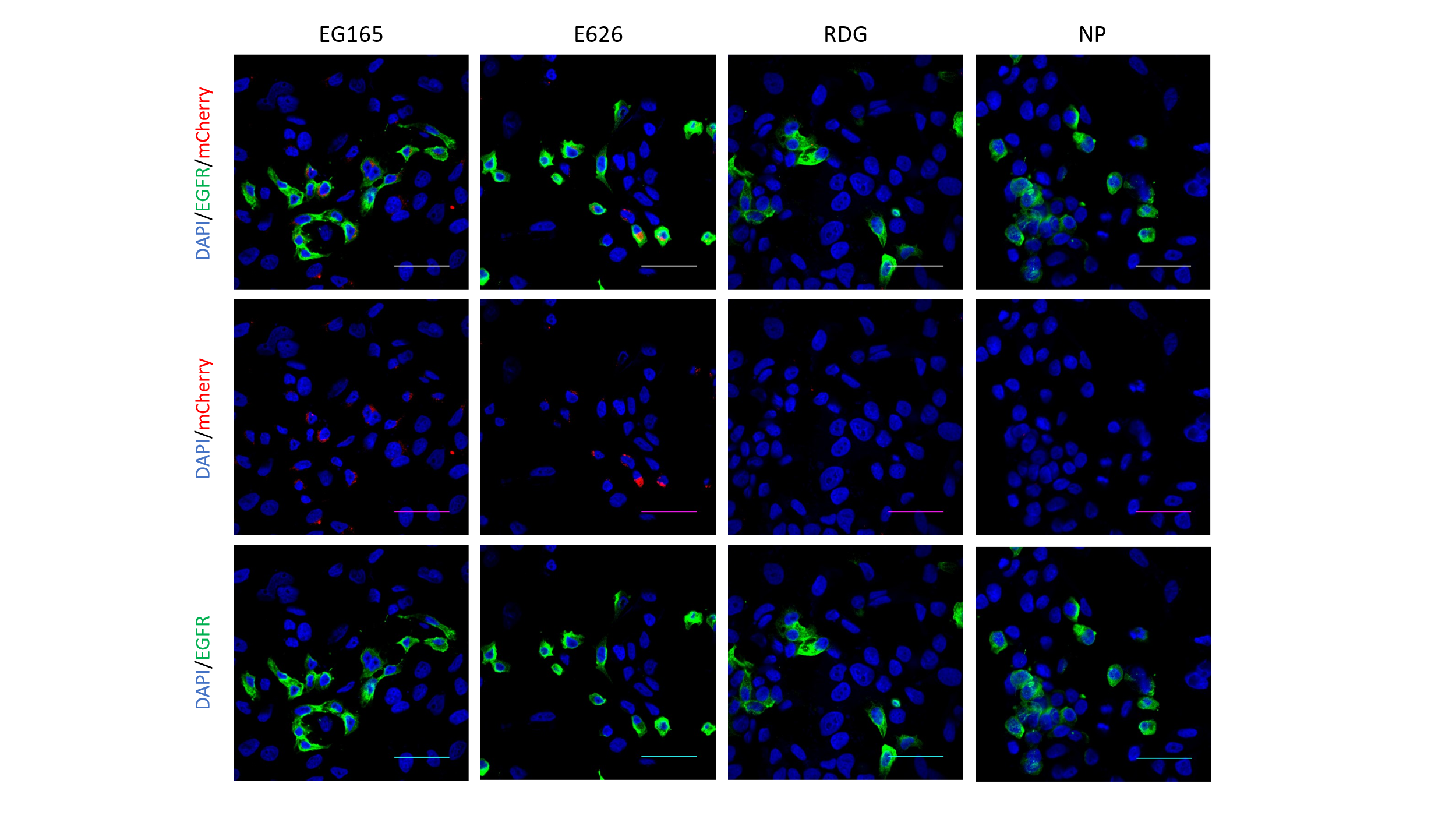

Figure S5. A431 and MCF-7 cells were co-cultured and treated with Monobody-mCherry co-labeled EVs for 10 min. The cells were fixed, and the binding was assessed by confocal laser scanning imaging of EVs (red), anti-EGFR antibody (green), and nuclear staining with DAPI (blue). Scale bar is 50 µm.

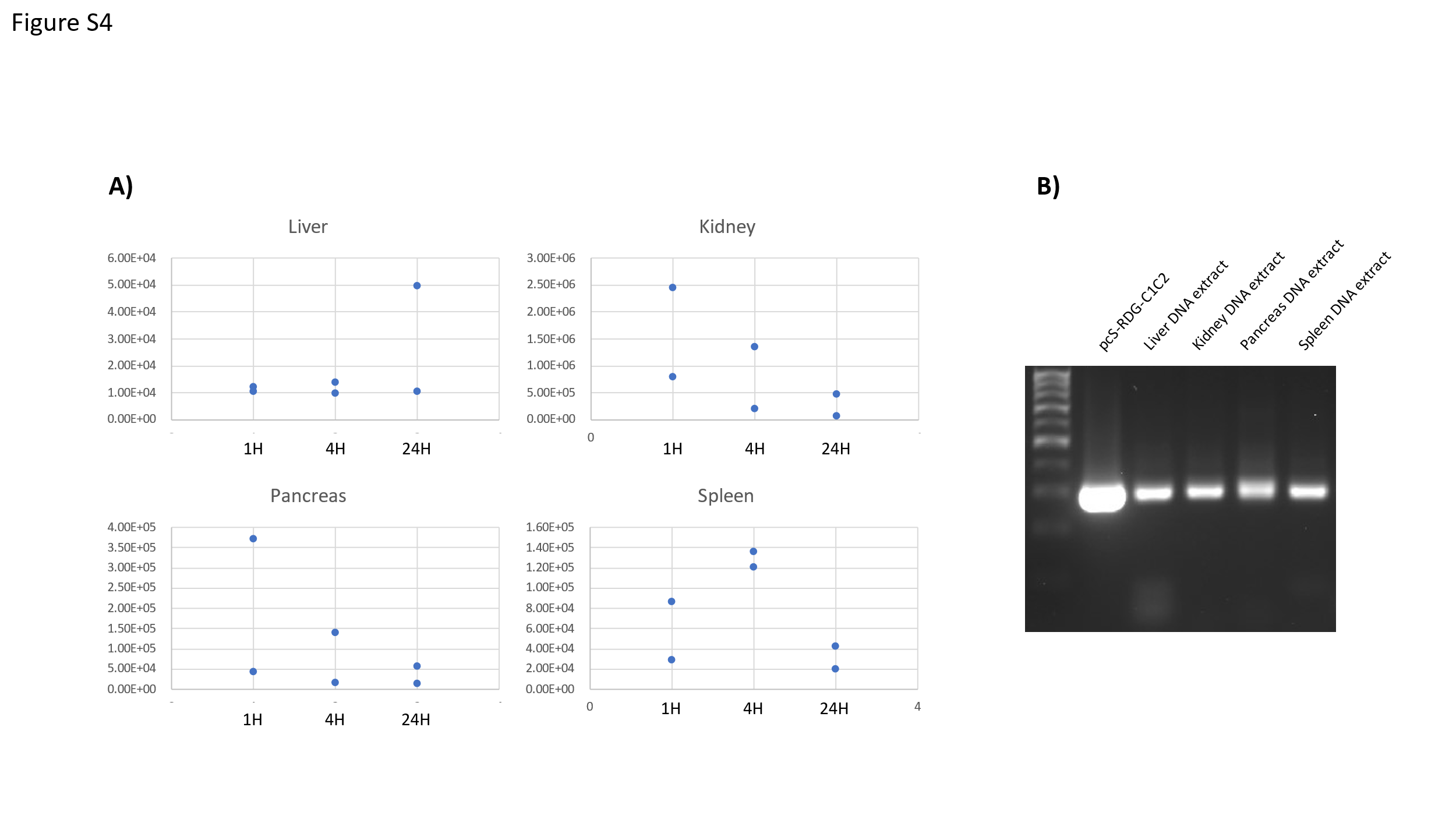

Figure S6. Time course evaluation of pDNA recovery from organ samples. Organ dissection was performed at 1H, 4H, and 24H post EV injection. (A)DNA amount from Liver, Kindey, Pancreas, and spleen was quantified by qPCR. (B)Amplified PCR products were analyzed on a gel to verify amplicon size.

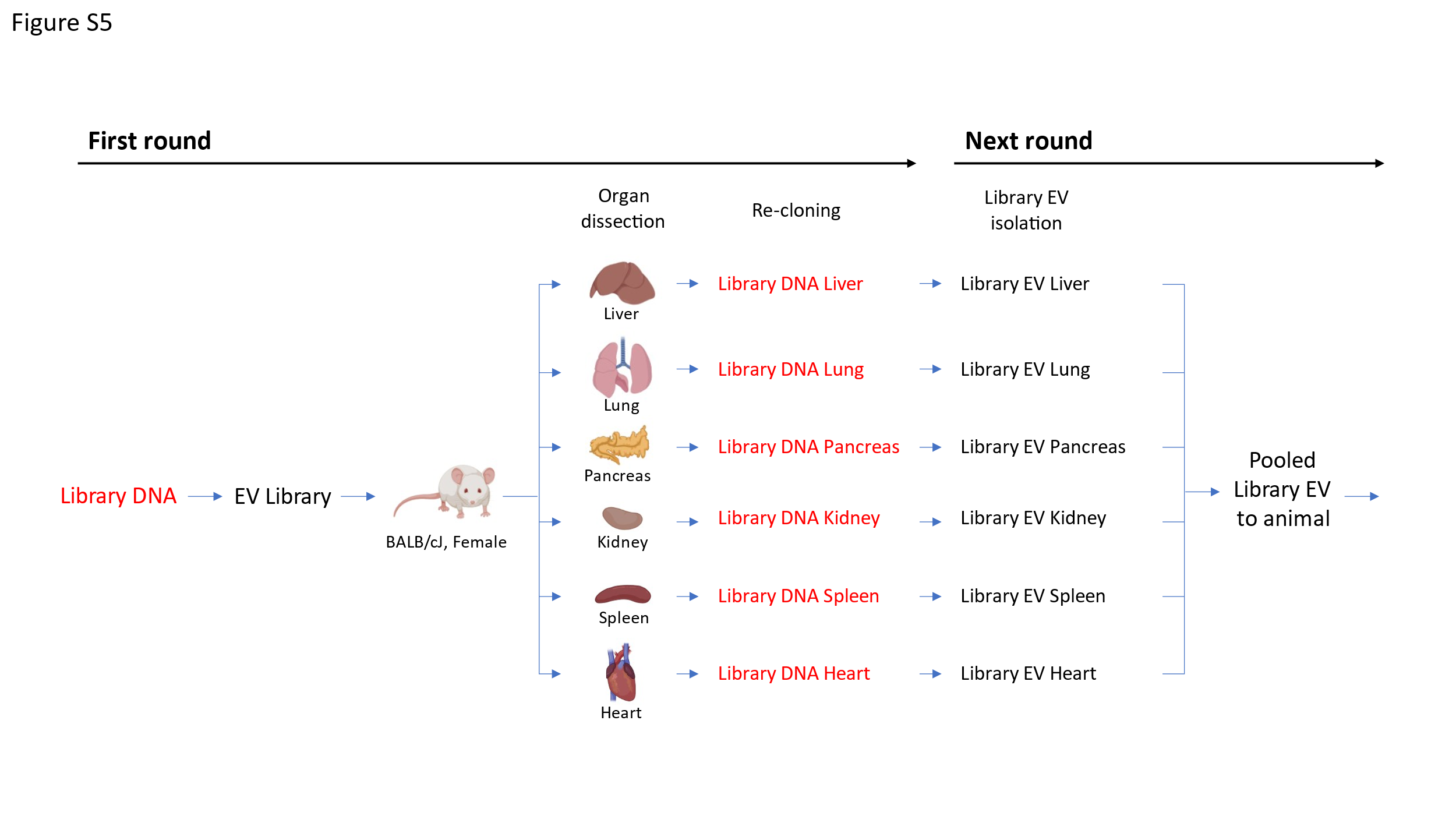

Figure S7. Schematic figure of in vivo EV library screening. EVs Library are injected to an animal via tail vein injection. After 1 hour circulation, organs (Heart, Liver, Lung, Kidney, Spleen, and Pancreas) are dissected followed by pDNA extraction. Extracted pDNA from each organ are re-cloned to generate Library DNA, followed by Library EVs generation. All organ specific EV Libraries are pooled and injected to an animal for next round of screening.

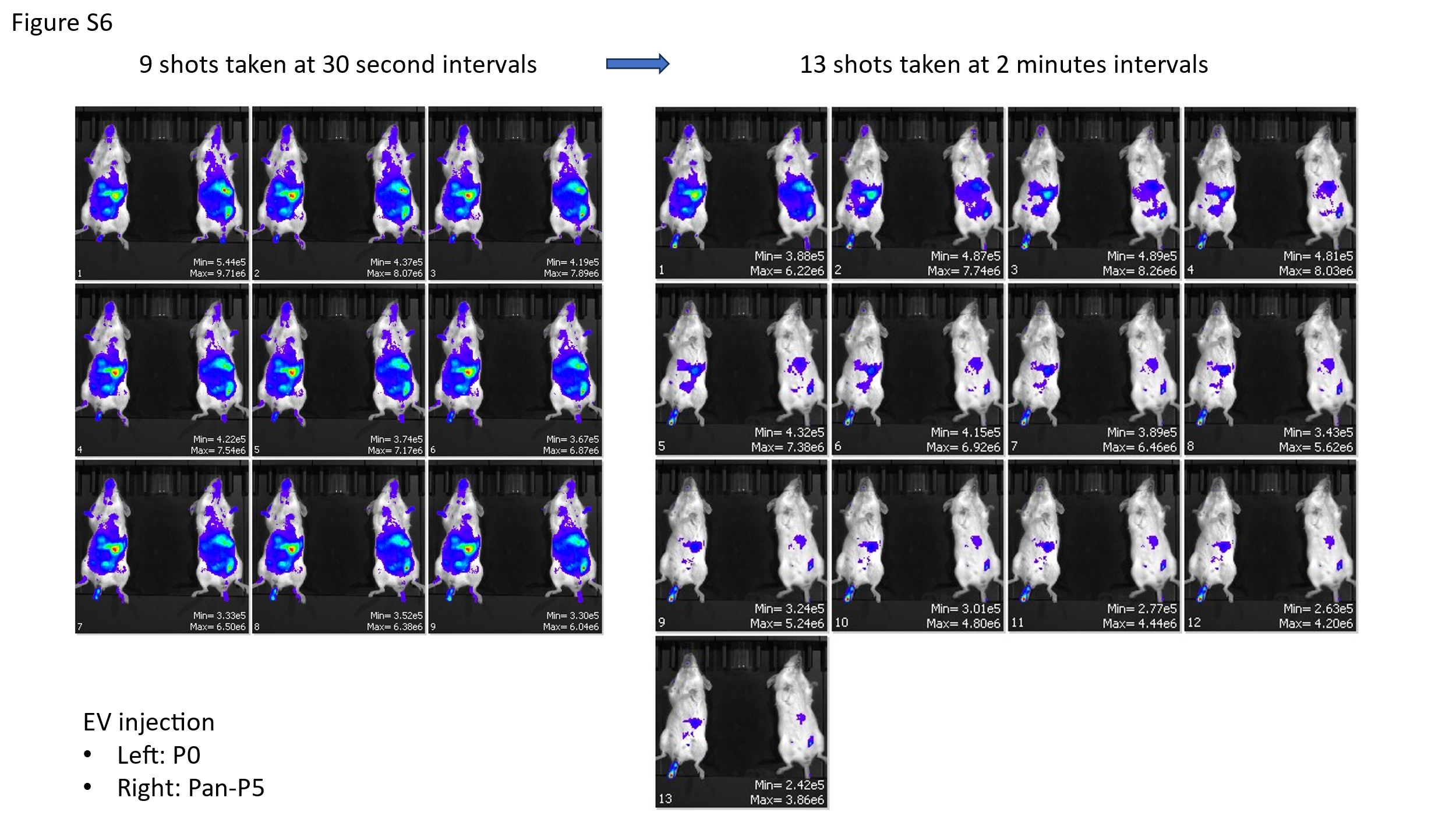

Figure S8. EV Library were introduced via tail vein injection. in vivo image was taken sequentially for 30 minutes to confirm distribution of NanoLuc labeled EVs Library.

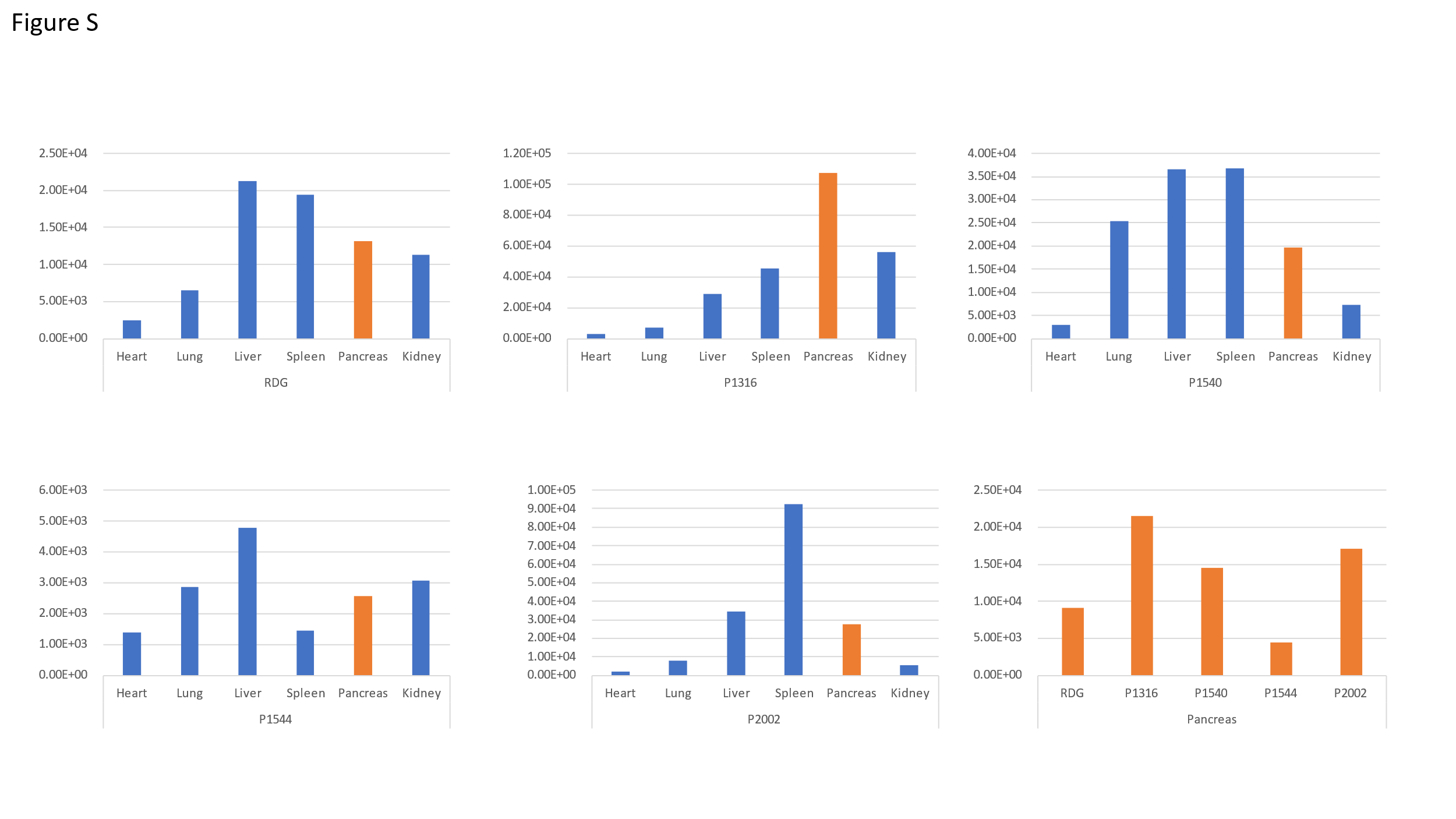

Figure S9. Preliminary evaluation of high binder candidates. Each monobody-EVs were co-labeled with NanoLuc, and introduced into mice. After 1H circulation, each organs were dissected and imaged by IVIS.

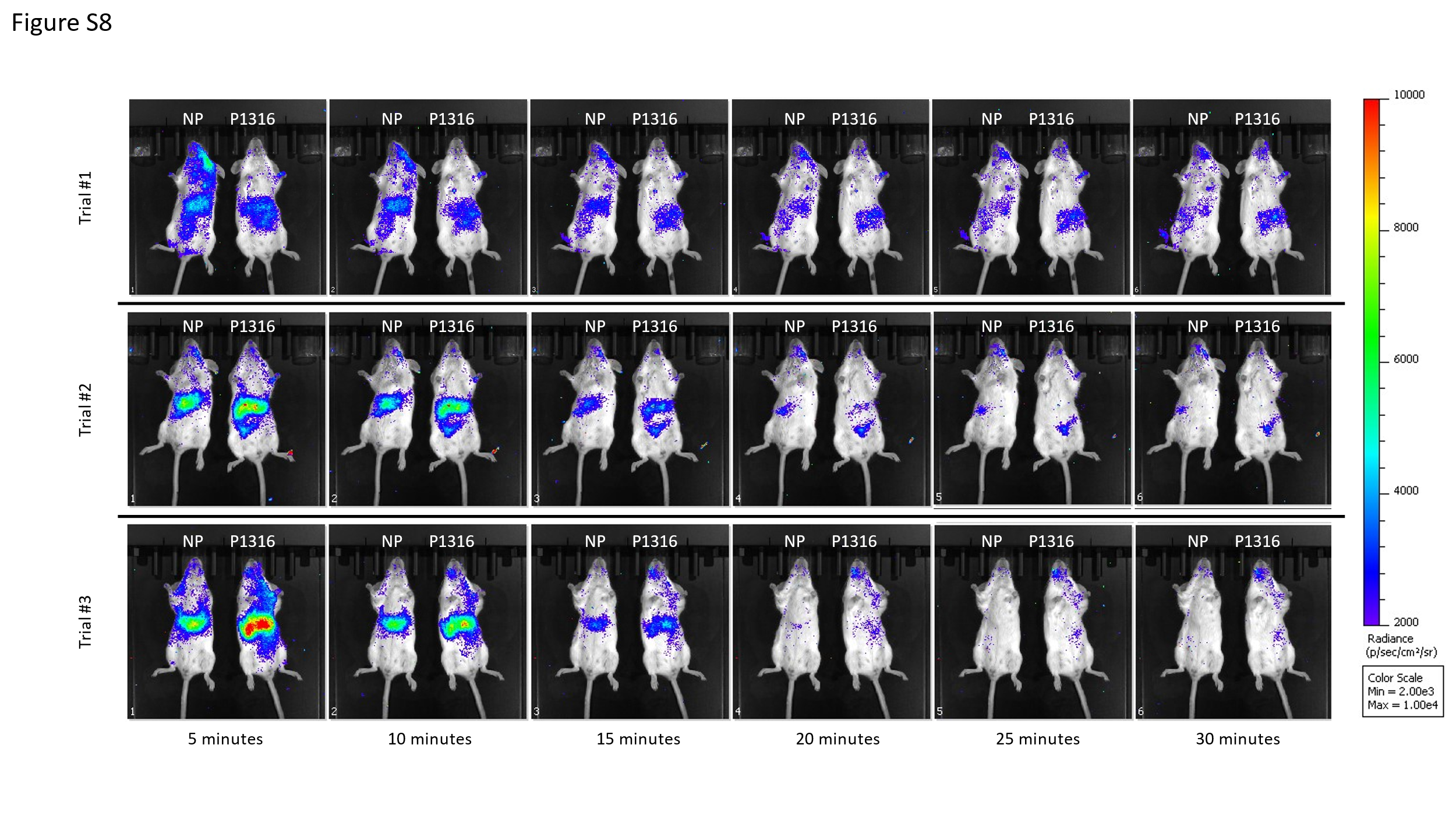

Figure S10. In vivo imaging of EV delivery to cells. ThermoLuc-CD63 co-labeled P1316 EVs were delivered via intravenous injection and biodistribution was monitored every 5 min up to 30 min.
